# Core and rod structures of a thermophilic cyanobacterial light-harvesting phycobilisome

**DOI:** 10.1101/2021.11.07.467602

**Authors:** Keisuke Kawakami, Tasuku Hamaguchi, Yuu Hirose, Daisuke Kosumi, Makoto Miyata, Nobuo Kamiya, Koji Yonekura

## Abstract

Cyanobacteria, glaucophytes, and rhodophytes utilize giant, light-harvesting phycobilisomes (PBSs) for capturing solar energy and conveying it to photosynthetic reaction centers. PBSs are compositionally and structurally diverse, and exceedingly complex, all of which pose a challenge for a comprehensive understanding of their function. To date, three detailed architectures of PBSs by cryo-electron microscopy (cryo-EM) have been described: a hemiellipsoidal type, a block-type from rhodophytes, and a cyanobacterial hemidiscoidal-type. Here, we report cryo-EM structures of a pentacylindrical allophycocyanin core and phycocyanin-containing rod of a thermophilic cyanobacterial hemidiscoidal PBS. The structures define the spatial arrangement of protein subunits and chromophores, crucial for deciphering the energy transfer mechanism. They reveal how the pentacylindrical core is formed, identify key interactions between linker proteins and the bilin chromophores, and indicate pathways for unidirectional energy transfer.

## Introduction

Cyanobacteria, glaucophytes, and rhodophytes utilize a large water-soluble, light-harvesting complex called phycobilisome (PBS) for solar energy absorption and energy transfer to photosynthetic membrane proteins (photosystem I and photosystem II; PSI and PSII)^1^. PBSs absorb light in the wavelength range 490–650 nm (as an exception, cyanobacteria containing chlorophyll f have PBSs that absorb near-infrared light^2^, which is otherwise difficult for PSI and PSII to harness (Extended Data Fig. 1). PBS is composed of phycobiliproteins (PBPs) such as phycoerythrin, phycoerythrocyanin, phycocyanin (PC), and allophycocyanin (APC). Assembly units of the PBPs are α- and β-subunits that have globin folds and harbor several linear tetrapyrrole chromophores such as phycoerythrobilin, phycourobilin, phycoviolobilin, and phycocyanobilin (PCB)^3^. Oligomers of these α- and β-subunits are assembled through non-chromophorylated linker proteins to construct the whole PBS complex. Five types of PBS structural morphology have been reported: hemidiscoidal^4,5^, hemiellipsoidal^6^, block-type^7^ rod-type^8,9^, and bundle-type^10^. Hemidiscoidal PBSs have bicylindrical^11^ tricylindrical^4^, or pentacylindrical cores^5^. Composition of PBPs with respect to number of rods, the overall structure of the PBSs, and their association with PSI and PSII are regulated by availability of light and nutrients^9, 12^. Recently, the three-dimensional (3D) structures of three types of red algal PBSs were analyzed by cryo-electron microscopy (cryo-EM)^13–16^, and details of their tricylindrical cores and energy transfer pathways were revealed. Since PBSs are compositionally and structurally diverse and exceedingly complex, structural and functional analyses of PBSs in different species are essential for comprehensive understanding of their working mechanisms.

The PBS of *Thermosynechococcus vulcanus* NIES-2134 (hereafter referred to as *T. vulcanus*) has a hemidiscoidal structure with a pentacylindrical APC core and six PC rods, the total molecular weight of which reaches a few MDa or more^16, 17^ (Extended Data Fig. 2). These are composed of α- (CpcA, ApcA, ApcD, and ApcE) and β- (CpcB, ApcB, and ApcF) subunits, and their basic constituent unit is an αβ monomer^18^. PCBs are covalently bound via a thio-ether linkage to a conserved cysteine residue in the α and β subunits^19^. Three PC monomers (trimer, (αβ)_3_) paired with two trimers form a disk (hexamer, (αβ)_6_), and two disks interact with each other to form a PC rod; the arrangement and number of PC rods vary among species^4, 5, 9^. Linker proteins in the PC rods are believed to contribute not only to structural stability between the disks but also to adjustment of the energy level of the chromophores. Linker proteins of *T. vulcanus* are classified as rod linker (L_R_; CpcC); rod-terminal linker (L_RT_; CpcD); rod–core linkers (L_RC_; CpcG1, CpcG2, and CpcG4); core–membrane linkers (L_CM_; ApcE), which bind the core to thylakoid membranes; or core linkers (L_C_; ApcC), which bind to the PBS core. When the PBS absorbs light, the excitation energy is transferred at a very fast rate (of the order of picoseconds) to chromophores in subunits at the membrane surface called terminal emitters (L_CM_, ApcD, and ApcF) and then eventually to PSII and PSI^20^. Several X-ray crystal structures of PCs and APCs from *T. vulcanus* have been reported, and these have provided starting points for discussion around possible overall structural configurations and mechanisms of internal energy transfer in PBSs^21–27^.

Here, we report the structures of the pentacylindrical APC core and PC rod from *T. vulcanus* at resolutions of 3.7 Å and 4.2 Å, respectively. The structures reveal the detailed architecture of hemidiscoidal PBSs and indicate a possible mechanism for the unidirectional excited energy transfer. They provide a basis for the understanding of the many cyanobacterial PBSs and could be utilized for applications for the efficient use of solar energy^27–29^.

## Results and Discussion

### Structural refinement and overall structure of PBS core

The pentacylindrical APC core (PBS core) was purified from preparations of PBSs followed by gradient fixation (GraFix) treatment^17, 31^. Most PC rods were absent probably because of dissociation during preparation of the cryo-EM grids (see Materials and Methods). Cryo-EM maps of the PBS core and PC rod were reconstructed to 3.7 Å and 4.2 Å resolutions, respectively, based on the Gold Standard Fourier shell correlation [FSC] criteria of 0.143 between two half maps (Extended Data Table 1). The structural models were refined against the maps, which validated the resolutions; FSC of 0.5 between the model and the map: 3.8 Å (PBS core) and 4.2 Å (PC rod); Q-score^32^ based estimates: 3.6 Å (Q = 0.49; PBS core) and 4.0 Å (Q = 0.40; PC rod) (Extended Data Figs. 3 and 4, and Extended Data Tables 1–3).

The PBS core of *T. vulcanus* shows a hemidiscoidal structure with *C*2 symmetry, composed of three cylinders (A, A’, and B), two cylinders (C and C’), and PC rods (Fig. 1). The strict twofold symmetry was not held in the outer part of the PC rods (Rb, Rb’, Rt, Rt’, Rs1, Rs1’, Rs2, and Rs2’)^17^, as many of these parts likely dissociated from the core during sample preparation (see above). The cryo-EM map resolves portions of the rods, but local resolution of the corresponding regions ranges from 7 to 20 Å (Extended Data Fig. 3). Thus, we did not build models of the rods (Fig. 1C). The PBS core is a supercomplex with the dimensions 110 × 210 × 300 Å^3^ and composed of 38 ApcAs, 40 ApcBs, 6 ApcCs, 2 ApcDs, 2 L_CMs_, 2 ApcFs, and 84 PCBs (Extended Data Fig. 5, Extended Data Table 4). The number of APC trimers in each cylinder of the PBS core differs between species^*4, 5, 13, 14, 16, 17*)^, implying that the architecture of a PBS affects the efficiency of energy transfer within the PBS as well as from PBS to PSII/PSI.

**Fig. 1.**
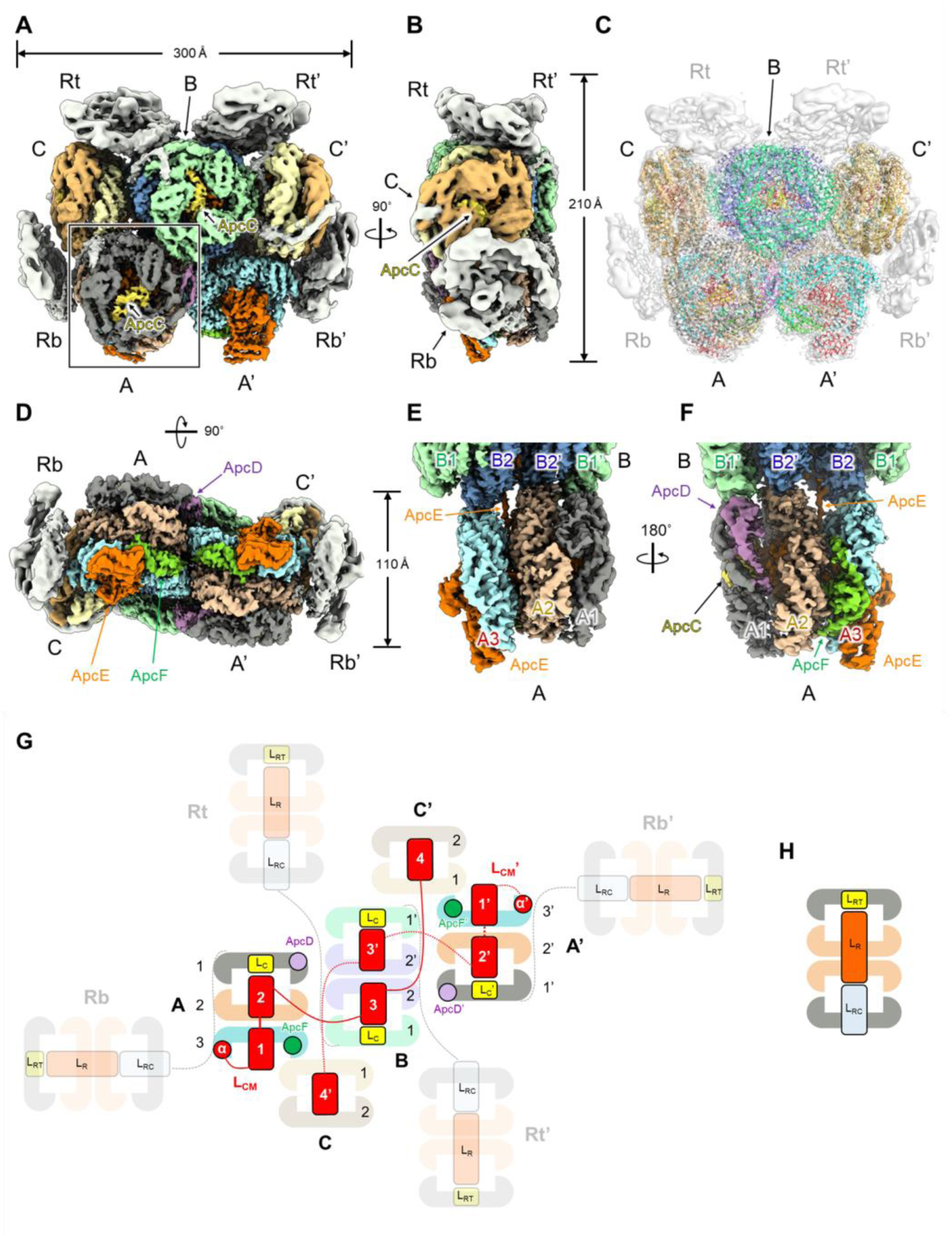
Overall structure of the PBS core from *T. vulcanus*. **(A-F)** Cryo-EM density map of the PBS core from the front (A, C), side (B, E, and F), and bottom (D) views. (C) shows the superposition of the cryo-EM density map and the refined PBS core model. (E) shows the structure inside the rectangle in (A), enlarged and rotated 90°. (F) shows (E) rotated 180°. **(G)** Schematic model of the pentacylindrical APC core (including A (A’), B, and C (C’) cylinders) and PC rods (Rb, Rb’, Rt, and Rt’). ApcE (L_CM_) containing α, Reps 1*–*4 and Arms 1*–*3 is shown in red. PC rods that did not build the model are shown in translucent. The PC rod models were drawn referencing the cryo-EM map in this study and the PBS structure from *Anabaena* sp. PCC 7120^*5*^. **(H)** Schematic model of the PC rod. CpcC (L_R_), CpcD (L_RT_), and CpcG (L_RC_) interact within the two PC hexamers.

The A cylinder is composed of the α-subunits of phycobiliproteins (ApcA, ApcD, and L_CM_), the β-subunits of phycobiliproteins (ApcB and ApcF), and ApcC (Lc). The A (A’) cylinder consists of three APC trimers (A1, A2, and A3) with subunits of ApcD/ApcB and two ApcA/ApcB (A1); three ApcA/ApcB (A2); and ApcE/ApcB, ApcA/ApcF, and ApcA/ApcB (A3), respectively. Electrophoretic analysis shows that ApcD is present in prepared PBSs (Extended Data Fig. 2)^17^, but ApcD could not be identified in the cryo-EM density map due to the limited resolution. Then, we tentatively assigned ApcD to a subunit that could not be identified as ApcA (Extended data Fig. 6). The B cylinder consists of four APC trimers consisting of ApcA and ApcB (B1, B2, B1’, and B2’). In addition, L_C_ interacts with one side of the A and C cylinders, while it interacts with both sides of the B cylinder. The C cylinder consists of two APC trimers, ApcA and ApcB, which correspond to a half B cylinder.

L_CM_ is a terminal emitter that is located at the membrane surface in the A cylinder and transfers energy to PSII. L_CM_ is composed of α^LCM^, showing a similar structure to ApcA, and has four repeated motifs named “repeats” (Rep1–Rep4) and “arms” (Arm1–3) that connect the motifs. Rep1 interacts with ApcF, two ApcAs (α2 and α3), and one ApcB (β1) in A1 (Fig. 2B).

**Fig. 2.**
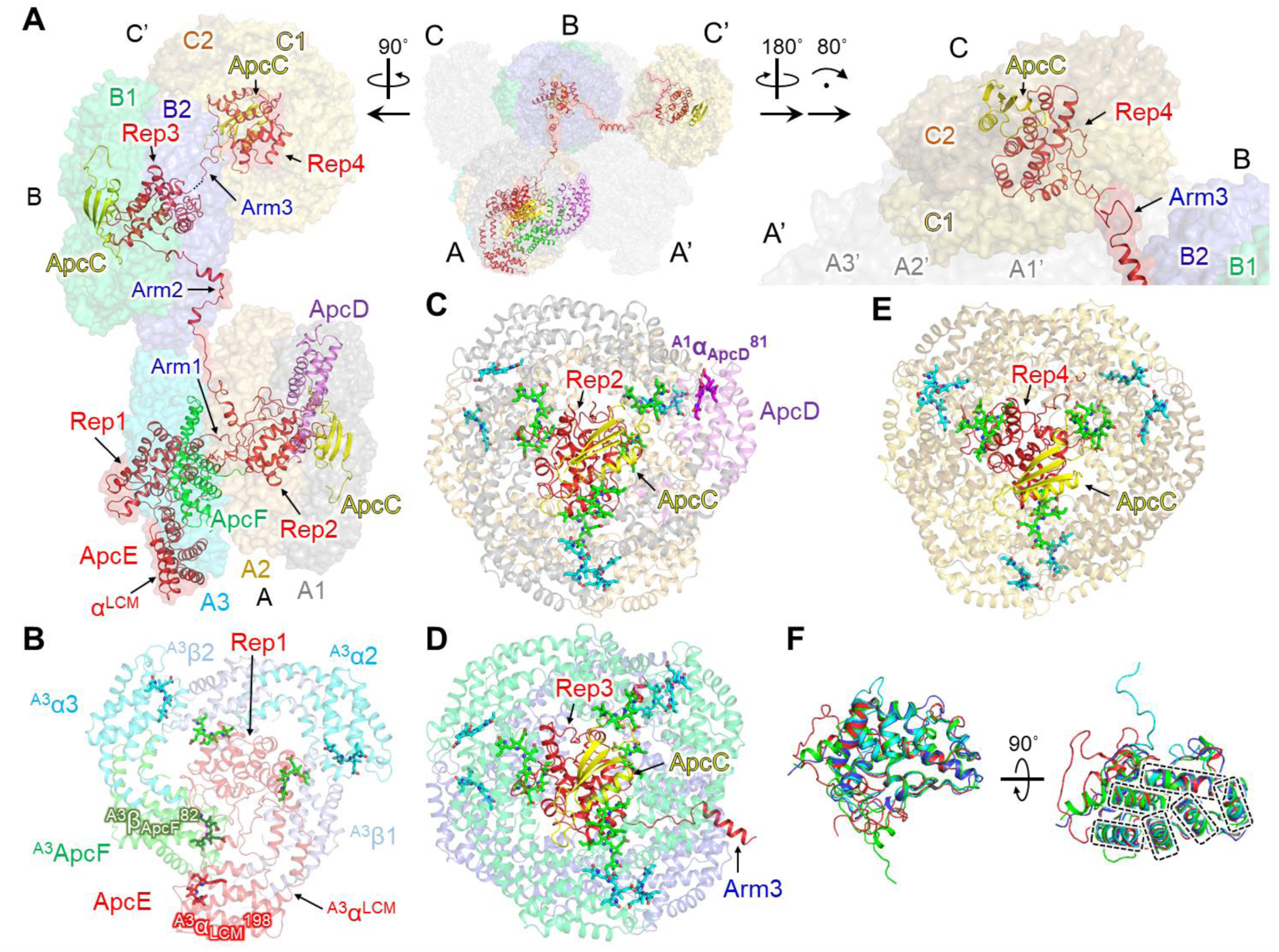
Structures of A, B, and C cylinders, including terminal emitters and linker proteins. **(A)** Arrangement of the terminal emitters (α^LCM^ of L_CM_, ApcD, and ApcF) and linker proteins (L_CM_ and ApcCs). **(B)** Structures of APC trimer (A3) and Rep1 of L_CM_ in A cylinder. **(C)** Structures of APC trimers A1 and A2, ApcC, and Rep2 of L_CM_ in A cylinder. **(D)** Structures of APC trimers B1 and B2, ApcC, Rep3, and Arm3 of L_CM_ in B cylinder. **(E)** Structure of APC trimers C1 and C2, ApcC, and Rep4 of L_CM_ in C’ cylinder. **(F)** Superposition of four Rep structures (Reps1–4). Dashed lines indicate the α-helixes in the Rep regions. Rep1, green; Rep2, red; Rep3, blue; Rep4, cyan.

In the recently reported structures of red algae, ApcC interacts with Rep1 of L_CM_, whereas ApcC and ApcD interact with Rep2 of the PBS core of *T. vulcanus*, indicating that the interaction between subunits in the core also differs between species. Rep3 of L_CM_, together with ApcC, contribute to the maintenance of the structure and function of the two APC trimers in the B cylinder. Arm3 extends to Rep4, which serves as the linker domain of the C cylinder. Rep4 interacts with ApcC in the same way as Rep2 and Rep3, which means that each cylinder in the PBS core contains one ApcC (Figs. 2C-E). An interesting feature of Reps1–4 in L_CM_ is that the structures of the α-helix regions are similar, whereas the structures of the loop regions differ (Fig. 2F). Sequence homology among Reps is not high (Extended Data Fig. 7), suggesting not only that Reps1–4 maintain the structure of each cylinder but also that distinct amino acid residues in each Rep around PCB could contribute to efficient transfer of absorbed light energy to the terminal emitters. Arm3, Rep4 of L_CM_, and the C cylinder are not present in the PBS core of red algae^13, 14^ or *Synechocystis* sp. PCC 6803^4^, whereas these components are present in about half of cyanobacteria if the L_CM_ domains are mapped to the phylogenetic tree of major cyanobacterial species (Extended Data Figs. 8 and 9). Thus, common to many species of algae including cyanobacteria, the C cylinder, Rep4, and Arm3, which maintain the C cylinder itself, must play a key role in light-harvesting. L_CM_ is one of the most important linker proteins for energy transfer to PSII. The structure of *T. vulcanus* PBS described in this study provides a starting point towards understanding component architecture and organizational networks for general working mechanisms of PBSs in the broad algae group.

### Arrangement of chromatophores in the PBS core and energy transfer pathway

The excitation energy is transferred to the terminal emitters (L_CM_, ApcD, and ApcF) via chromophores in the B and C (C’) cylinders, and the energy is eventually transferred to PSII and PSI. *T. vulcanus* has PCB as its only chromophore, and it is the protein environment around each chromophore that must be intimately involved in the transfer of the excited energy to the terminal emitters. Indeed, amino acid residues with polar/charged groups affect the absorption energy of pigments^33^. In addition, an asparagine residue in the β subunits is methylated (ligand ID: MEN), and its mutation prevents the growth of cyanobacteria under high light conditions, indicating its importance in energy transfer^34^.

Figure 3 shows each cylinder that constitutes the PBS core, the chromophores in the core, and the distances between chromophores. In general, the excitation energy is transferred between close pairs of chromophores, i.e., the donor (D) and acceptor (A) (Förster Resonance Energy Transfer)^35, 36^. There are four main factors associated with excitation energy transfer rate: (i) the distance between D and A; (ii) the orientation factor, κ^2^, which is a factor that describes the relative orientation of the transition dipoles of D and A in space (Extended data Fig. 10); (iii) the overlap integral between the fluorescence and absorption spectra of D and A, respectively; and (iv) quantum yield of D in the absence of A. For a freely rotating pigment D and A, κ^2^ has a mean value of 2/3, and the values of κ^2^ the significant chromophores are approximately 1–3 (Extended Data Table 5, Extended Data Figs. 10 and 11). This indicates that energy transfer is likely to occur between their chromophores in the PBS core. The efficiency is inversely proportional to the sixth power of the distance between D and A through dipole–dipole coupling; thus, the shorter the distance between chromophores, the greater is the efficiency of the energy transfer between them. Therefore, energy transfer efficiency between cylinders is strongly also dependent on the distance between close chromophores.

**Fig. 3.**
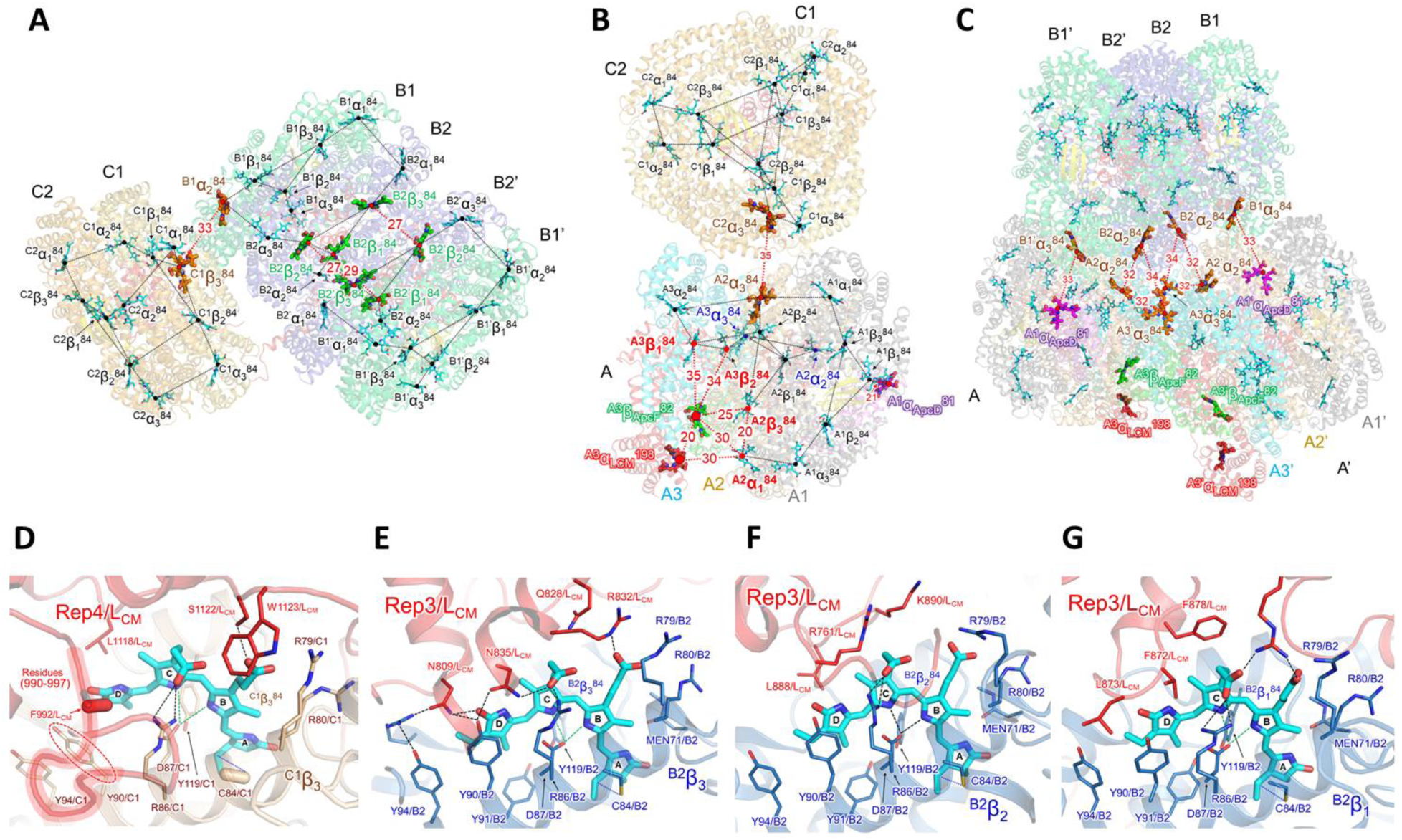
Arrangement of chromophores is the key to the energy transfer pathway in PBS core. **(A)** Possible energy transfer pathways in and between B and C (C’) cylinders. **(B)** Possible energy transfer pathways in and between A and C cylinders. **(C)** Possible energy transfer pathways between A (A’) and B cylinders. **(D-G)** The chromophores and the surrounding amino acid residues of L_CM_. The numbers near the dotted lines indicate the distances (Å) between the PCB pairs. In **(E-G)**, dashed lines (black and green) indicate hydrogen bonds, and dotted lines (blue) indicate covalent bonds between PCB and cysteine residue. In the amino acid residues (Phe992/L_CM_ and Cys84/C1) in **(D)**, the directions of atoms from C_α_ to C_β_ in the residues are indicated by the capsule-shaped objects.

The chromophores in the PBS core are numbered according to the nomenclature of those in red alga^13, 14^, and their names are defined as follows: cylinder name (A, B, or C), subunit composition (α or β), and the number of cysteine residues (84) to which the chromophore is bound (Extended Data Fig. 12). Again we reiterate that the previous red algal PBS structures do not contain C cylinders^13, 14^. The chromophores involved in the energy transfer between or within each cylinder are shown in Fig. 3A-C. The energy transfer from the C (C’) cylinder to the B and A (A’) cylinders occurs via ^C1^α_1_^84^–^B1^α_2_^84^ (34 Å) and ^C2^α_3_^84^–^A2^α_3_^84^ (35 Å), respectively (Fig. 3A-B). In the present structure, the D ring of PCB in many ApcBs forms π–π interactions with tyrosine residues (Y90), but the D ring of ^C1^β_3_^84^ likely interacts with Phe992 within the loop region (residues 990-997) of Rep4 in the L_CM_ (Fig. 3D). In addition, ^C1^β_3_^84^ interacts with S1122/L_CM_. These interactions probably contribute to energy transfer from the C (C’) cylinder to the B cylinder. In the cyanobacterial PBS with bound PC rods (Rt, Rb, Rs1, and Rs2)^5, 17^, the energy received by the PC rods is believed to be transferred to the B and A (A’) cylinders via the C (C’) cylinder, indicating that the C (C’) cylinder acts as an intermediary for energy transfer from the PC rods to the terminal emitters.

The B cylinder consists of four APC trimers, with two APC trimers (B2 and B2’) interacting with each other. This indicates that energy transfer in the B cylinder is mediated by ^B2^β_2_^84^–B^2’^β_1_^84^ (27 Å), ^B2^β_1_^84^–^B2’^β_1_^84^ (29 Å), and ^B2^β_3_^84^–^B2’^β_3_^84^ (27 Å) (Fig. 3A). Energy transfer from the B cylinder to the A (A’) cylinder seems to involve ^B1’^α_3_^84^–^A1^α_ApcD_ ^81^ (33 Å) and ^B2^α_2_^84^–^A3’^α_3_^84^ (34 Å) (Fig. 3C). In addition, there is an interaction between the A and A’ cylinders, suggesting energy transfer occurs through ^A2^α_2_^84^–^A3’^α_3_^84^ (32 Å). These chromophores interact with Rep3 and Rep4 of L_CM_, and the amino acid residue compositions of L_CM_ differ around each of the four chromophores (^C1^β_3_^84, B2^β_3_^84, B2^β_2_^84^, and ^B2^β_1_^84^) (Fig. 3D-G). Three chromophores (^B2^β_3_^84, B2^β_2_^84^, and ^B2^β_1_^84^) interact with amino acid residues in Rep3/L_CM_, significantly altering the peripheral structure of each chromophore. ^B2^β_3_^84^ forms hydrogen bonds with asparagine residues (N809/L_CM_ and N835/L_CM_), and basic amino acids (R762/L_CM_ and K890/L_CM_) are present near ^B2^β2 ^84. B2^β3^84^ interacts with a basic residue (R730/L_CM_) and is surrounded by hydrophobic amino acids (L873/L_CM_, T872/L_CM_, and F878/L_CM_). This difference in the protein environment around each chromophore may define their individual energy level.

### Excited energy transfer to terminal emitters

Three PCBs in the A cylinder (^A2^α_3_ ^84, A2^α2^84^, and ^A3^α_3_^84^) are believed to be the key chromophores that receive excitation energy from the B and C cylinders based on their spatial arrangement (Fig. 3B), which then ultimately pass it to the two chromatophores (^A3^β_ApcF_ ^82^ and ^A3^α_LCM_ ^198^) in the terminal emitters. In this case, the excitation energy is most likely transferred via the four chromophores (^A2^α_1_^84, A2^β_3_^84, A3^β1^84^, and ^A3^β_2_^84^) close to these two chromophores. Three of the four chromophores interact with Rep1 or Rep2, in a characteristic structure (Fig. 4A-C). The κ^2^ value and the distance between chromophores also suggest that the energy transfer efficiency of ^A3^α_LCM_^198^–^A3^β_ApcF_^82^ is highest among all the identified pairs (Extended Data Table 5).

^A3^β_1_^84^ interacts with Y306/L_CM_ and is surrounded by aromatic amino acids (Y428/L_CM_ and F432/L_CM_). ^A3^β_2_^84^ interacts with S348/L_CM_ and R366/L_CM_, and F373/L_CM_ is close to the D ring of ^A3^β2^84^. In addition, ^A2^β_3_^84^ interacts with R630/LCM and is surrounded by three aromatic amino acids (Y455/L_CM_, Y610/L_CM_, and F637/L_CM_), with F637/L_CM_ close to the D ring of ^A2^β_3_^84^. It was noted that, in the structure of red algal PBSs, aromatic amino acids are abundant in the vicinity of the chromophores and it was proposed that they regulate the energy state of each chromophore to achieve unidirectional energy transfer^13,14^. The aromatic amino acids in the linker proteins of *T. vulcanus* PBS may function similarly, suggesting that the protein environment around the three chromophores is optimized for efficient transfer of excitation energy to the two chromophores.

The chromophore ^A3^β_ApcF_ ^82^ in ApcF binds C82/ApcF as well as C416/ L_CM_ and interacts with four amino acid residues (Y265/ L_CM_, R84/ApcF, R77/ApcF, and R78/ApcF). In the absence of both ApcF and ApcD, the PBS is unable to transfer energy to either PSII or PSI, and thus the major route of energy transfer to PSII and PSI appears to be through ApcF rather than ApcD^37^. Thus, the protein environment around ^A3^β_ApcF_ ^82^ in ApcF seems key for transferring the unidirectional excitation energy to PSII and PSI. ApcD is a terminal emitter that transfers energy to PSI^37^, and the present study suggests that ^A1^β_1_^84^ and ^B1^α_3_^84^ are involved in the energy transfer pathway to ^A1^α_ApcD_ ^81^.

We were not able to identify the side chains of three amino acid residues (W166/L_CM_, D163/L_CM_, and K159/L_CM_), as the cryo-EM density map around ^A3^α_LCM_^198^ was somewhat disordered (Fig. 4E, Extended Data Fig. 3E). Nevertheless, the present structure suggests that ^A3^α_LCM_ ^198^ in L_CM_ could form a π–π interaction with W166/L_CM_, as the D-ring of PCB and tyrosine residues in ApcA commonly form a π–π interaction. This feature is probably an important factor for energy transfer from ^A3^α_LCM_ ^198^ to PSII. Further details with regard to ^A3^α_LCM_ ^198^ and key interactions with surrounding structures will need a higher-resolution structure of the PBS and/or a supercomplex structure interacting with PBS and PSII.

**Fig. 4.**
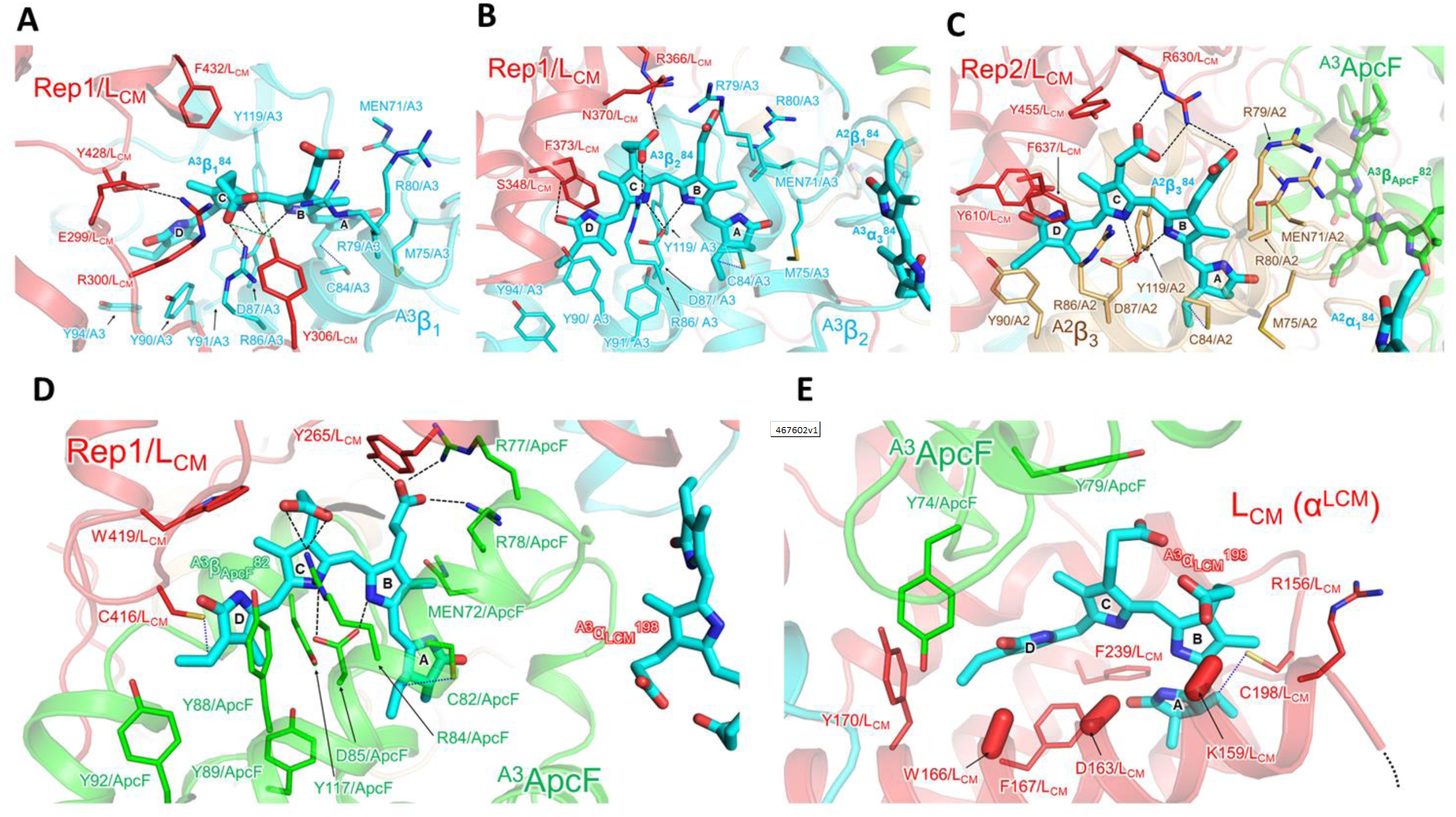
Chromophores and their surrounding structures in terminal emitters (ApcF and L_CM_). **(A-B)** Chromophores in the APC trimer A3 and their interaction with Rep1 of L_CM_. **(C)** Chromophore in the APC trimer A2 and its interaction with Rep2 of L_CM_. **(D)** Chromophore of ApcF and its interaction with Rep1 of L_CM_. **(E)** Chromophore of L_CM_ and its interaction with ApcF. The dashed lines (black) indicate hydrogen bonds. The dotted lines (blue) indicate covalent bonds between PCB and cysteine residue. The directions of atoms from Cα to Cβ in the residues are indicated by the capsule-shaped objects.

### Structure of PC rod and arrangement of chromatophores

The PC rod forms a pentadecamer with two PC hexamers (Disks A and B) interacting endwise with each other (Fig. 5A) and three linker proteins inside the rod. CpcC, CpcD, CpcG1, CpcG2, and CpcG4 have been identified as linker proteins of the PC in *T. vulcanus* PBSs (Extended Data Fig. 2)^17^, and we could assign CpcC, CpcD, and CpcG2 in the cryo-EM map of the PC rod (Fig. 5B-C). As CpcG1, CpcG2, and CpcG4 show similar amino acid sequences (Extended Data Fig. 13), CpcG2 assignment in this study is tentative.

**Fig. 5.**
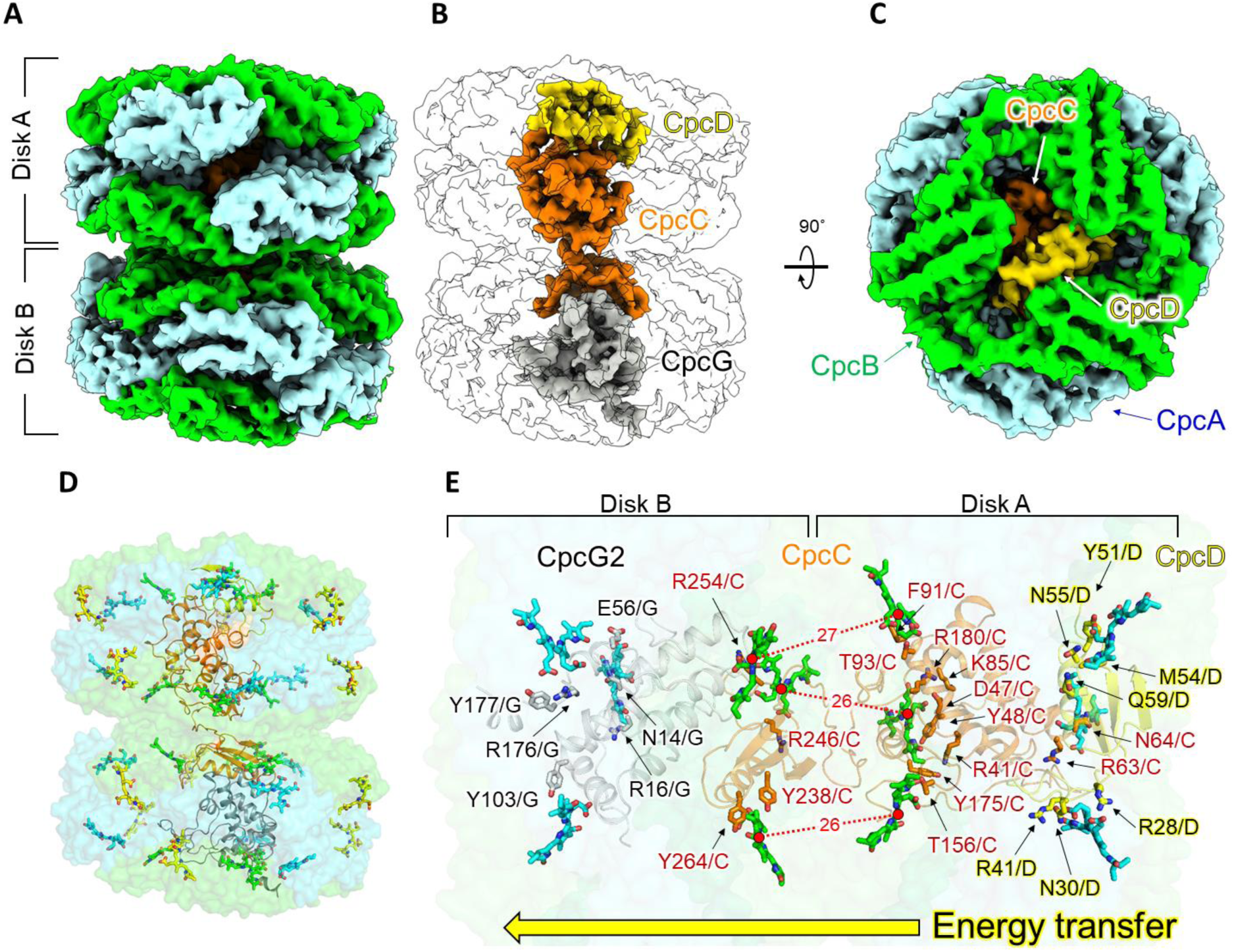
Structure of PC rod, including linker proteins, and its energy transfer pathway. **(A)** cryo-EM density map of the PC rod (Disk A and Disk B). **(B)** Arrangement of the linker proteins (CpcC, CpcD, and CpcG) in the PC rod. **(C)** Structure of the PC rod in (B) rotated 90°. **(D)** Arrangement of chromophores in the PC rod. α^84^, β^84^, and β^155^ are colored cyan, green, and yellow, respectively. **(E)** Arrangement of β^84^s in a PC rod interacting with linker proteins. β^84^s at the boundary between Disk A and Disk B are colored green, and these chromophores are involved in the energy transfer between Disk A and Disk B. The amino acid residues of the linker proteins near β^84^s are indicated in (E).

The PC rods dissociated from the PBS should contain different CpcG subunits. In the *Anabaena* PBS, when three CpcG encoding genes (*CpcG1, CpcG2*, and *CpcG4*) were deleted, the PC rods could not bind to the PBS core^5^. When one of the CpcG encoding genes was deleted, the PC rods could interact with the PBS core but not in the same manner as the wild-type PBS. This suggests that the composition of the CpcG protein is different in each PC rod^5^. The cryo-EM density map resolves the linker proteins extending from the PC rods interacting with the PBS core, with a resolution enough to see (Fig. 1, Extended Data Fig. 14). The C-terminal region of the CpcG protein in Rt (Rt’) and Rb (Rb’) mainly interacts with the peripheral region of the B cylinder and the peripheral region of the A (A’) cylinder, respectively. The C-terminal region of CpcG interacts weakly with the cylinders of the PBS core and also supports a rod.

The PC structures obtained by X-ray crystal structure analysis do not include any linker proteins, and the interaction between the PC monomers in the oligomer is symmetric^9^. In contrast, the interaction between PCs and linker proteins in the PC rod is asymmetric (Fig. 5A and D), which is induced by the interactions between CpcC, CpcD, and CpcG2 in the rod. Consequently, the arrangement of chromophores, especially at the boundary between Disks A and B in the PC rod, is different from that inferred from the crystal packing of the PC structure (Fig. 5D, Extended Data Fig. 15). CpcD (L_R_), a small rod cap linker domain, is the outermost linker protein of the intact PBS, and the excitation energy is transferred from Disk A to Disk B in the PC rod via the chromophores (β84s) interacting with CpcC (distances between chromophores are 26–27 Å) (Fig. 5E). A significant difference is noted in the chromophore arrangement in the cryo-EM map, especially in Disk B, although the observed distances were almost the same as those predicted from the crystal structure analysis. This difference is apparently caused by interaction of linker proteins (especially CpcC and CpcG2) (Extended Data Fig. 15). In previous studies on PC using spectroscopy, β84 in a PC rod interacting with linker proteins was proposed to be involved in the energy transfer between PC hexamers^*38*–*40*^. Therefore, six β84s (Fig. 5E) at the boundary between Disks A and B are probably involved in the energy transfer between the disks. The characteristic amino acid residues around the chromophores in the linker proteins and the interactions between these residues and the chromophores are likely crucial for unidirectional energy transfer in the PC rod.

## Methods

### Preparation of intact PBSs, PBS cores, and PC rods

Cells of the thermophilic cyanobacteria *Thermosynechococcus vulcanus* NIES-2134 were cultured in a phosphate medium^40, 41^. Intact PBSs and PBS cores were prepared according to the protocols described by Kawakami et al.^17^. Thylakoid membranes, PSI, and PSII were prepared according to the protocols described by Kawakami et al.^40, 41^. For PC rods, the intact PBSs were mounted on a holey carbon film-coated copper grid (see **“Sample preparation and data collection for cryo-EM”** for a description of pre-treatment of the grid), and then a small amount of buffer containing 10 mM phosphate buffer was quickly added to reduce the phosphate concentration in the sample to less than 0.5 M. This resulted in the dissociation of the PC rods from the intact PBSs.

### Sample preparation and data collection for cryo-EM

To clarify the structure of the PBS core and PC rod, grids for cryo-EM were prepared in the following manner. A holey carbon film-coated copper grid (Quantifoil R1.2/1.3 Cu 200 mesh, Microtools, GmbH, Berlin, Germany) that had been pretreated by Au sputtering was glow-discharged for 10 s using an ion coater (JEC-3000FC, JEOL, Tokyo, Japan). For the PBS core, 3.0 μL of the PBS core was applied to the grid and blotted with filter paper for 4 s, then immediately plunge-frozen in cooled ethane using a FEI Vitrobot Mark IV (Thermo Fisher Scientific, Waltham, MA, USA) under 100% humidity at 4 °C. For the PC rod, 3.5 μL of the purified sample was applied to the grid and diluted with 1.5 μL of 10 mM phosphate buffer on the grid. Then, the grid was blotted and manually plunge-frozen in cooled ethane using a homemade plunger. The grids were then introduced into a CRYO ARM 300 electron microscope (JEOL) equipped with a cold-field emission gun and an in-column energy filter with a slit width of 20 eV. Dose-fractionated images were recorded using a K2 summit direct electron detector (AMETEK, Berwyn, PA, USA) in counting mode. All images were corrected using a JEOL automatic data acquisition system^43^ with a nominal magnification of 40,000×, which corresponded to a pixel size of 1.24 Å. Nominal defocus ranges and dose rates for PBS core and PC rod were -0.5 to -1.5 μm and 84.1 e^-^ Å^-2^ with 50 frames, and -0.5 to -1.5 μm and 50.5 e^-^ Å^-2^ with 30 frames, respectively. In total, we collected 4,600 and 2,865 movies for the PBS core and PC rod, respectively.

### Data processing

The collected movie stacks of the PBS core were divided into eight optics groups to correct changes in the beam tilt over time. Drift correction and dose-weighted frame summing were performed using MotionCor2-1.3.2^44^, and contrast transfer function (CTF) parameters were estimated using CTFFIND4-1.10^45^. The images were selected for further data processing based on the Thon ring patterns. The PBS core particles were manually picked and subjected to reference-free, two-dimensional (2D) classification by RELION-3.1.0^46^ to create reference images for automatic particle picking. In all, 128,676 particles were picked automatically and extracted with a pixel size of 2.48 Å. Good averaged classes containing 45,427 particles after 2D classification were subjected to ab initio three-dimensional (3D) reconstruction in cryoSPARC-2.12.0^47^. Following the 3D classification in RELION, a well-aligned 3D class containing 25,532 particles was extracted with a pixel size of 1.24 Å and used for further 3D refinement with two-fold symmetry enforcement. Post-processing in RELION yielded a 4.75-Å resolution map based on the gold standard FSC. Finally, the resolution reached 3.71 Å after two rounds of Bayesian polishing and CTF refinement.

For PC12mer, all movie stacks were processed using MotionCor2-1.2.1 and Gctf-1.06^48^ for drift correction, frame summation, and CTF parameter estimation. The 812,327 particles were picked by convolutional neural network picking using EMAN-2.3.1^49^ and binned to a pixel size of 4.96 Å during extraction using RELION-3.0. After two rounds of 2D classification, 309,291 particles were selected and subjected to ab initio model construction using *cis*TEM-1.0.0 beta^50^. Good particles were automatically picked again with a 3D reference using the EMAN-2.3.1. In all, 159,537 particles with a pixel size of 1.86 Å were selected from 1,022,349 picked particles based on 2D and 3D classifications. 3D refinement using RELION-3.0 was performed, and following Bayesian polishing, CTF refinement and additional 3D classification improved the resolution to 4.19 Å with 111,054 particles without any symmetry enforcement. For further details, see Extended Data Figs. 4 and 5, and Extended Data Table 1.

### Model building, refinement, and validation

Initial models of the PBS core and PC rod from *T. vulcanus* were built using reference models of PC and APC (PDB codes: 3O18, 3DBJ) and homology modeling was performed on the SWISS-MODEL server (https://swissmodel.expasy.org/) by referring to the structures of the linker proteins (PDB codes: 5Y6P and 6KGX). The obtained models were fitted with the cryo-EM density maps using the “fit in map” program in UCSF Chimera (version 1.13), and initial models of the PBS core and PC rod were created. The initial models were modified manually using COOT to fit the cryo-EM density map and refined using Phenix (version 1.19.2). Subsequently, each subunit and its interacting subunits in the PBS core and PC rod were grouped and refined against the cryo-EM density map using REFMAC5 (version 5.8.0267) in CCP-EM. Finally, the grouped models were merged into one, and the overall structures (PBS core and PC rod) were refined using Phenix. The refinement statistics of the refined models were obtained using the comprehensive validation program in Phenix. The restraints needed for the ligands in the PBS core and PC rod from *T. vulcanus* were generated by electronic Ligand Bond Builder and Optimization Workbench (eLBOW)^51^. The restraint information for a methylated Asn (ligand ID: MEN) and phycocyanobilin (ligand ID: CYC) was obtained from the model of MEN identified in a high-resolution crystal structure (PDB code: 3O18). Comprehensive validation (in Phenix), Q-score^32^ and FSC-Q^52^ were used to validate the refined structural models (PBS core and PC rod).

### Estimation of the orientation factor between chromophores in the PBS core

Based on the arrangement of the PCBs and their orientation in the constructed PBS core, an approximate orientation factor, *κ*^2^, was estimated (Extended Data Table 5). These estimates were made especially for the sites likely to be involved in the energy transfer between cylinders in the PBS core and for the terminal emitters. Excitation energy transfer (EET) rate (*k*_EET_) is given by the following equation (1), where *V* and Θ are the electronic coupling factor and the overlap integral, respectively.

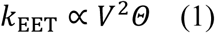

As shown in fig. S10 (A), the transition dipole moment of donor (D) and acceptor (A) are ***μ***_**D**_ and ***μ***_**A**_, respectively. **r** is the intermolecular center-to-center distance between D and A. *V* is written by the approximation equation as shown in the following equations (2, 3), and the orientation factor (*κ*^2^) can be estimated by equation (4).

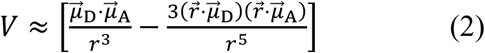

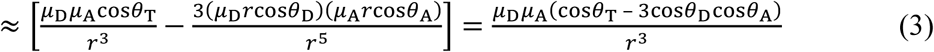

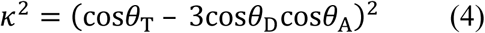

The orientation of the transition dipole moment of PCBs was determined by referring to Yang et al^53^ (Extended Data Figs. 10 and 11).

### Absorption spectroscopy

Absorption spectra of the thylakoid membrane, PSII, PSII, and PBS were measured at 23°C using a UV-2600 spectrophotometer (Shimadzu, Kyoto, Japan). The spectrum of the thylakoid membrane was measured using the opal-glass method.

### Polypeptide analysis and mass spectrometry

Polypeptides denatured with dithiothreitol were separated on a SuperSep gel containing 10–20% acrylamide (Wako, Tokyo, Japan) and then stained with Coomassie Blue (Extended Data Fig. 2). A gel imaging system (ChemiDoc XRS+system; Bio-Rad, Hercules, CA, USA) was used to photograph the stained gels. Separated polypeptides (a band containing ApcC and CpcD) were identified by mass spectrometry. The obtained protein bands were treated by in situ digestion using trypsin^54^. The fragmented peptides were analyzed by peptide mass fingerprinting (PMF) and MS/MS using Autoflex speed (Bruker Daltonics). The obtained mass spectra were analyzed by MASCOT server (MATRIX SCIENCE).

### Phylogenic analysis

Thirty-eight sequences of ApcE from selected strains of cyanobacteria, glaucophytes, and rhodophytes were obtained by blastp program at NCBI website (https://blast.ncbi.nlm.nih.gov/Blast.cgi) (Extended Data Figs. 8 and 9). The sequences were aligned using mafft ver. 7.478 with L-INS-i option^55^. The maximum likelihood tree of the ApcE sequences were estimated using iqtree2 ver. 2.1.4 with LG+F+R5 model selected by ModelFinder. The statistical support of the trees was estimated with 1000 replications of ultrafast bootstrap approximation^56^. Domain organization of ApcE was determined using hmmscan program in HMMER ver. 3.3.2 (http://hmmer.org/) with Pfam database^57^. Phylogenetic tree and domain organization were visualized using iToL ver. 4^58^.

## Acknowledgments

We thank research assistants Ms. Yuko Kageyama (Biostructural Mechanism Laboratory, RIKEN SPring-8 Center) and Ms. Rie Uno (Osaka City University) for their help with cell cultures, preparation of samples, electrophoresis analysis, and spectroscopies. We also thank Ms. Tomomi Shimonaka at the Graduate School of Science, Osaka City University, for performing MS/MS spectrometry.

## Author contributions

KK and NK designed the study; KK prepared the samples and performed electrophoresis analysis; KK measured spectroscopies, TH measured EM micrographs, and TH and KK processed the EM data (PBS core: TH, PC rod: KK) and reconstructed the final EM map (PBS core: TH, PC rod: KK); KK performed structural analysis; YH performed phylogenetic tree analysis; DK commented on data analyses; KY, NK, and MM supervised this project; KK, TH, and KY wrote the draft manuscript; KK, TH, and KY revised the final manuscript, and all of the authors contributed to the interpretation of the results and improvement of the manuscript.

## Data and materials availability

Atomic coordinates and cryo-EM maps for the reported structure of the PBS core and PC rod from *Thermosynechococcus vulcanus* were deposited in the Protein Data Bank under accession codes 7VEA (PBS core) and 7VEB (PC rod), and in the Electron Microscopy Data Bank under accession codes EMD-31944 (PBS core) and EMD-31945 (PC rod), respectively. Other data are available from the corresponding authors upon reasonable request.

