## Supplemental Figures and Tables for "Core and rod structures of a thermophilic cyanobacterial light-harvesting phycobilisome"

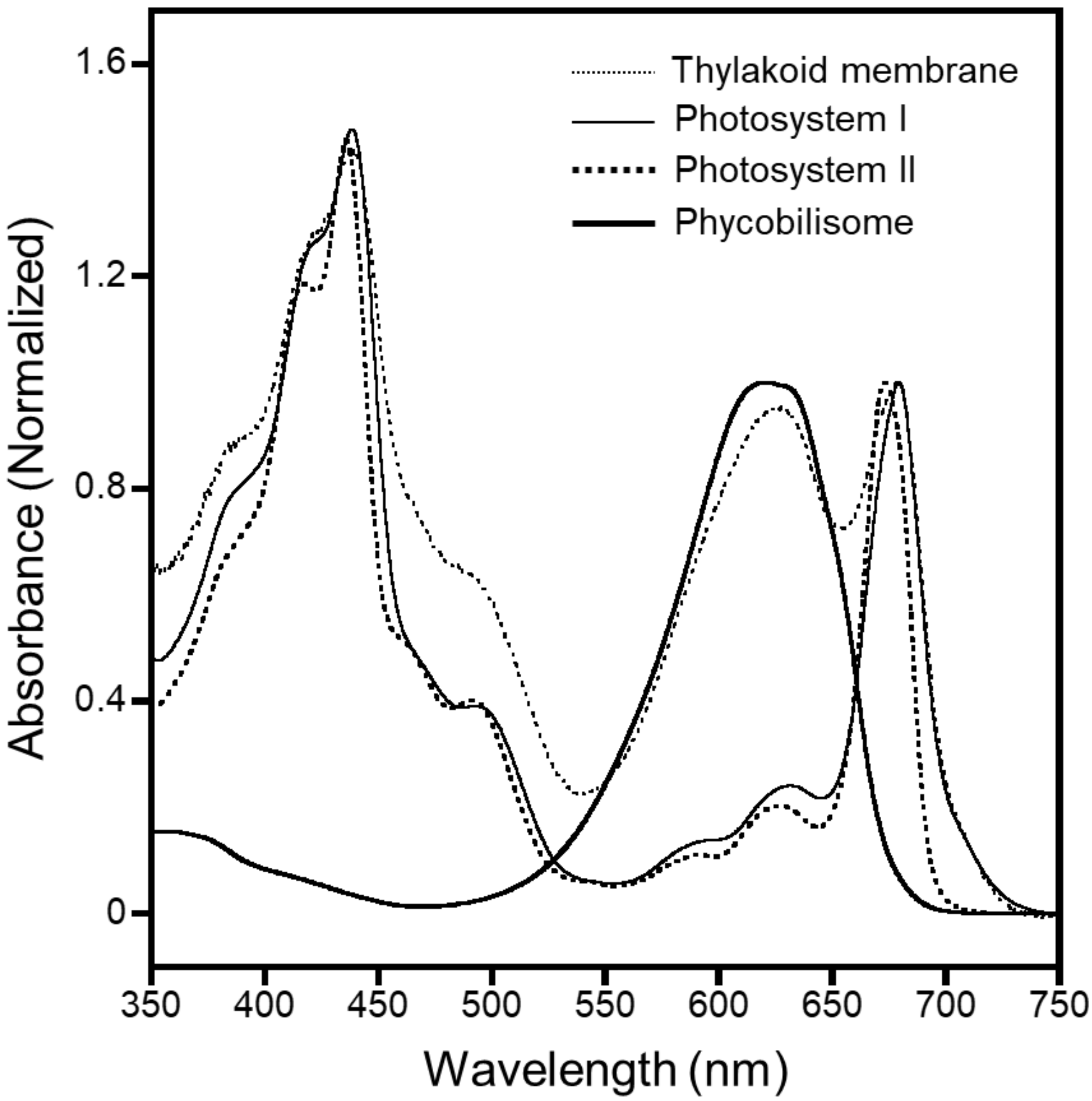


**Extended Data Figure 1. Absorption spectra of photosynthetic protein complexes.**

Thylakoid membrane, thin dotted line; photosystem I, thin solid line; photosystem II, bold dotted line; phycobilisome, bold solid line.

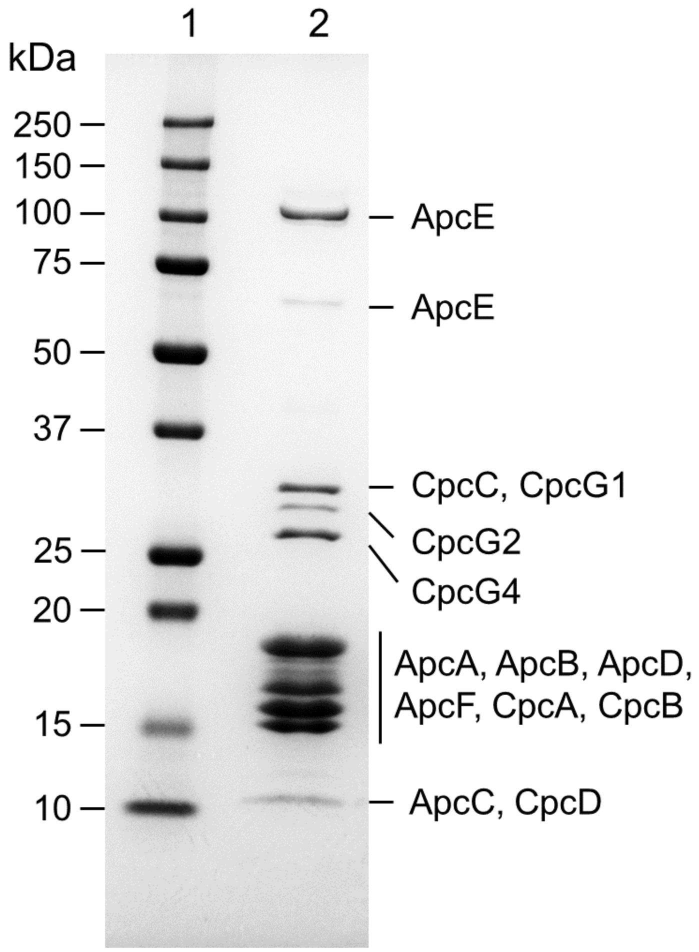


**Extended Data Figure 2. Polypeptide analysis.** lane 1, marker; lane 2, the prepared intact PBS.

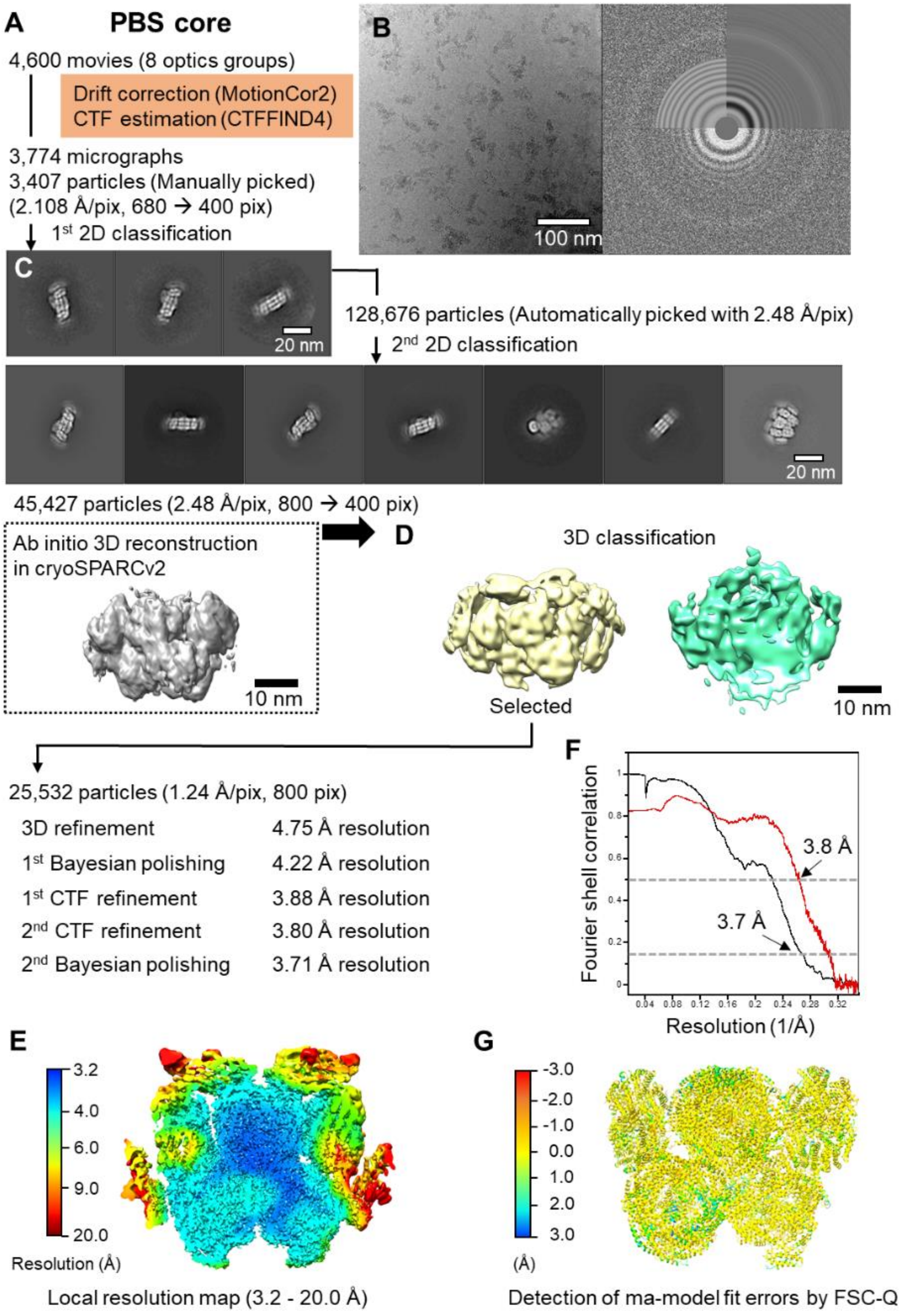
A
PBS core
4,600 movies (8 optics groups)
Drift correction (MotionCor2)
CTF estimation (CTFFIND4)
3,774 micrographs
3,407 particles (Manually picked)
(2.108 Å/pix, 680 → 400 pix)
1st 2D classification
B
100 nm
C
20 nm
128,676 particles (Automatically picked with 2.48 Å/pix)
2nd 2D classification
20 nm
45,427 particles (2.48 Å/pix, 800 → 400 pix)
Ab initio 3D reconstruction
in cryoSPARCv2
10 nm
D
3D classification
Selected
10 nm
25,532 particles (1.24 Å/pix, 800 pix)
3D refinement 4.75 Å resolution
1st Bayesian polishing 4.22 Å resolution
1st CTF refinement 3.88 Å resolution
2nd CTF refinement 3.80 Å resolution
2nd Bayesian polishing 3.71 Å resolution
F
Fourier shell correlation
3.8 Å
3.7 Å
Resolution (1/Å)
E
Resolution (Å)
Local resolution map (3.2 - 20.0 Å)
G
(Å)
Detection of ma-model fit errors by FSC-Q

**Extended Data Figure 3. Data collection and image processing workflow of the PBS core using cryo-EM. (A)** Overview of the data processing workflow. The resolution was estimated based on the gold standard Fourier shell correlation (FSC) criteria of 0.143. **(B)** Representative micrograph and Thon ring. The equivalent 4,600 images were used for data processing. **(C)** Good reference-free 2D class of the PBS core. **(D)** 3D classification of the PBS core. **(E)** A slice through the local resolution map of the PBS core to show the internal details. **(F)** FSC curves for 3D reconstruction of the cryo-EM map and the refined model versus the overall 3.7 Å map. Black, gold-standard curve with a value of 0.143 at 3.7 Å resolution; red, FSC curve calculated between the cryo-EM map and the refined structure model of the PBS core. The map-model FSC has a value of 0.5 at 3.8 Å resolution. **(G)** FSC-Q values calculated for the PBS core structural model. Atoms with FSC-Q values close to zero mean that the structural model is supported by the two half maps signal, while FSC-Q values that are positively away from zero corresponded to areas where the fit between the model and the map is low, or where the resolution of the map is low. On the other hand, negative FSC-Q values correspond to the atoms being correlated with noise, which means overfitting. Based on the FSC-Q validation, it can be assessed that the refined PBS core model is almost free of overfitting and that the model is reasonable for the cryo-EM map at 3.7 Å resolution.

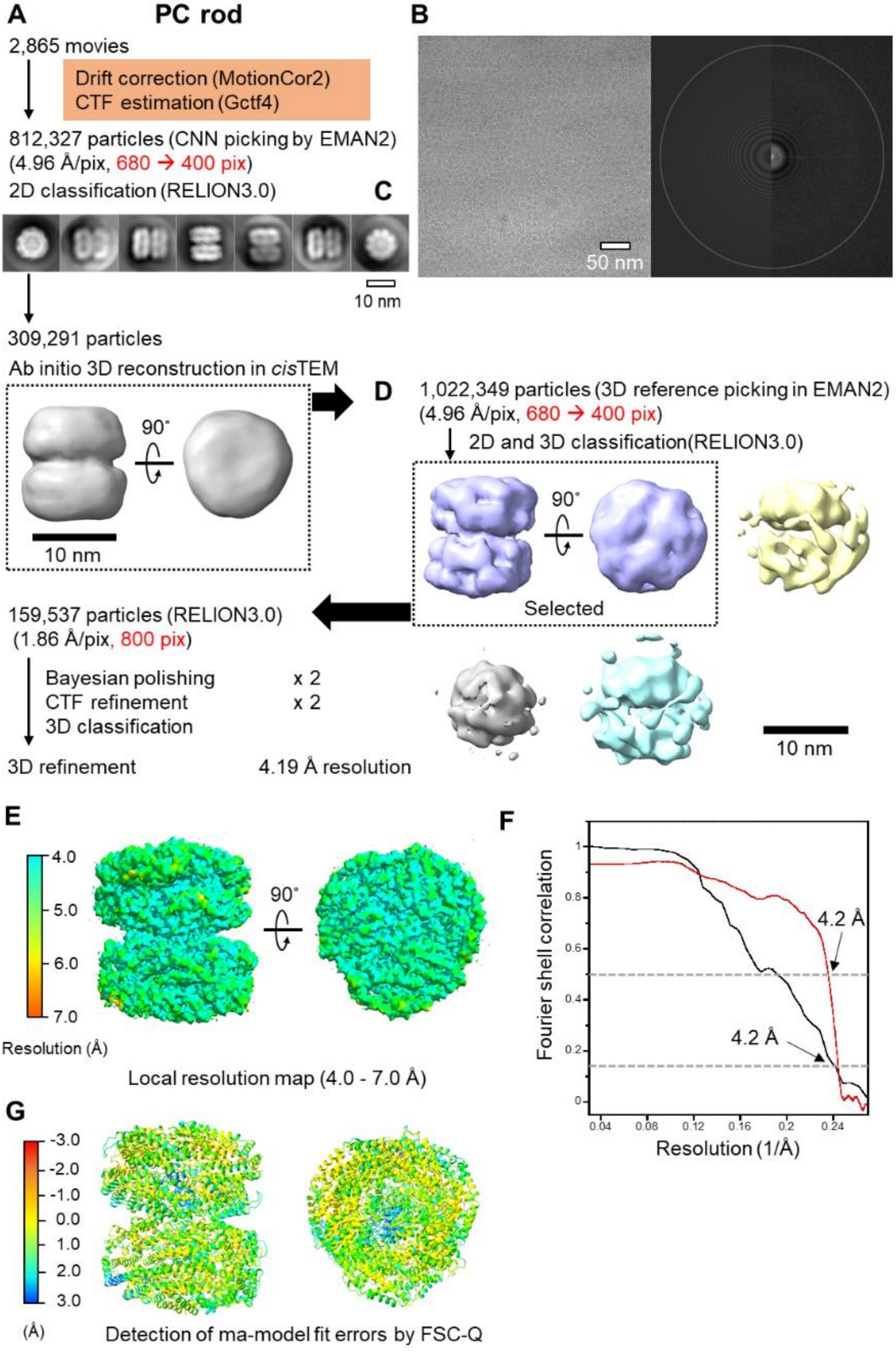

A
PC rod
2,865 movies
Drift correction (MotionCor2)
CTF estimation (Gctf4)
812,327 particles (CNN picking by EMAN2)
(4.96 Å/pix, 680 → 400 pix)
2D classification (RELION3.0)
C
10 nm
309,291 particles
Ab initio 3D reconstruction in cisTEM
90˚
10 nm
B
50 nm
D
1,022,349 particles (3D reference picking in EMAN2)
(4.96 Å/pix, 680 → 400 pix)
2D and 3D classification(RELION3.0)
90˚
Selected
10 nm
159,537 particles (RELION3.0)
(1.86 Å/pix, 800 pix)
Bayesian polishing x 2
CTF refinement x 2
3D classification
3D refinement
4.19 Å resolution
E
4.0
5.0
6.0
7.0
Resolution (Å)
90˚
Local resolution map (4.0 - 7.0 Å)
F
Fourier shell correlation
Resolution (1/Å)
4.2 Å
4.2 Å
G
-3.0
-2.0
-1.0
0.0
1.0
2.0
3.0
(Å)
Detection of ma-model fit errors by FSC-Q

**Extended Data Figure 4.**

**Data collection and image processing workflow of the PC rod by cryo-EM. (A)** Overview of the data processing workflow. The resolution was estimated based on the gold standard Fourier shell correlation (FSC) criteria of 0.143. **(B)** Representative micrograph and Thon ring. The equivalent 2,865 images were used for data processing. **(C)** Good reference-free 2D class of the PC rod. **(D)**3D classification of the PBS core. **(E)** The local resolution map of the PC rod. **(F)** FSC curves for 3D reconstruction of the cryo-EM map and the refined model versus the overall 4.2 Å map. Black, gold-standard curve with a value of 0.143 at 4.2 Å resolution; red, FSC curve calculated between the cryo-EM map and the refined structure model of the PC rod. The map-model FSC has a value of 0.5 at 4.2 Å resolution. (G) FSC-Q values calculated for the PC rod structural model. Atoms with FSC-Q values close to zero mean that the structural model is supported by the two half maps signal, while FSC-Q values that are positively away from zero corresponded to areas where the fit between the model and the map is low, or where the resolution of the map is low. In almost all cases for resolutions worse than 4 Å, the FSC-Q values of a structural model increase to >1.5 Å[47]. On the other hand, negative FSC-Q values correspond to the atoms being correlated with noise, which means overfitting. Based on the FSC-Q validation, it can be assessed that the refined PC rod model is almost free of overfitting and that the model is reasonable for the cryo-EM map at 4.2 Å resolution.

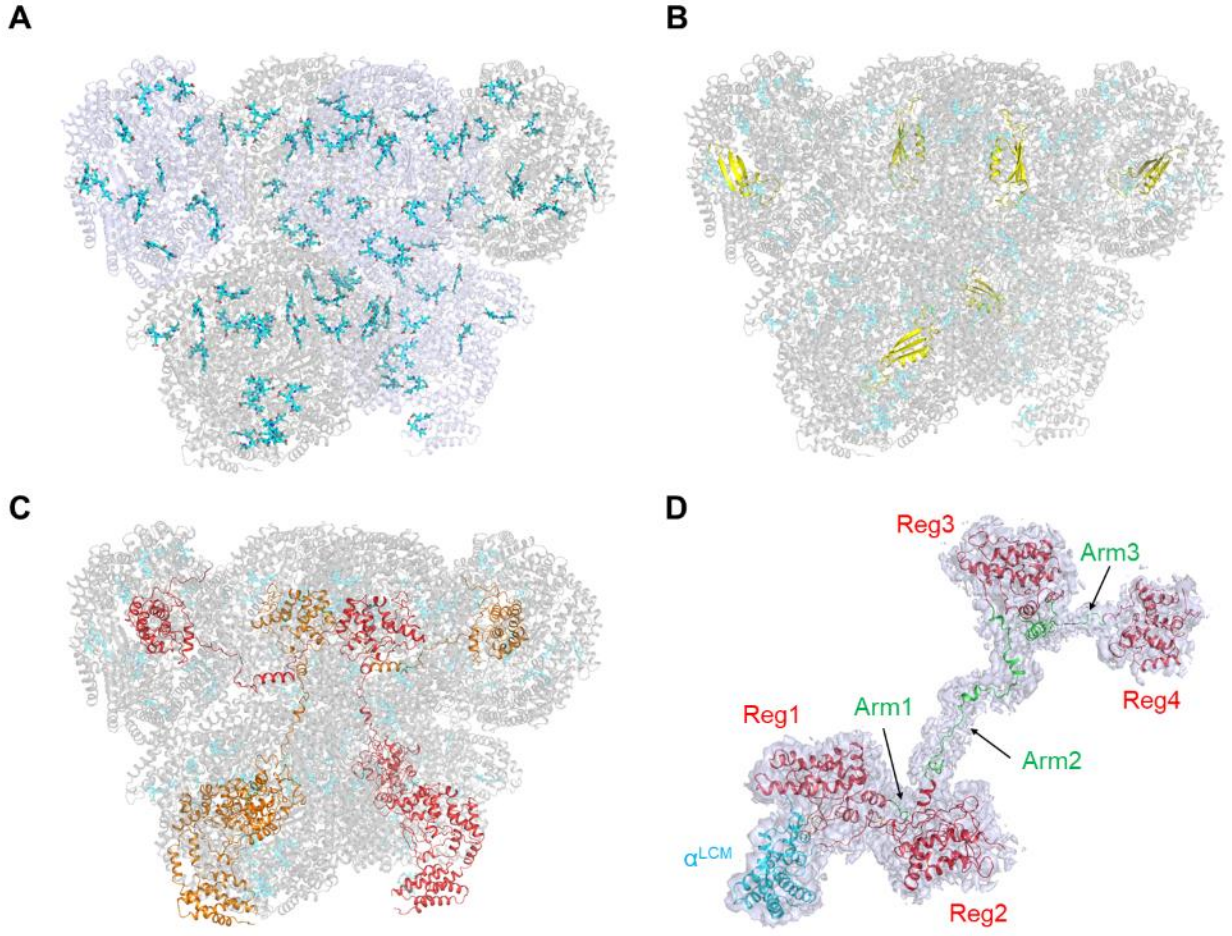


**Extended Data Figure 5. Arrangement of chromophores and linker proteins in the PBS core from *T. vulcanus*. (A)** Phycocyanobilins (cyan) distribution in the PBS core. **(B)** Arrangement of six ApcCs ($L_C$, yellow) in the PBS core. **(C)** Arrangement of two ApcEs ($L_{CM}$; orange and red) in the PBS core. **(D)** Cryo-EM map of ApcE ($L_{CM}$) and its refined model. ApcE is composed of the $\alpha^{LCM}$ (cyan), Reg1–4 (red), and Arm1–3 (green). Cryo-EM map of ApcE is shown in a surface representation at 1.0 sigma contour levels.

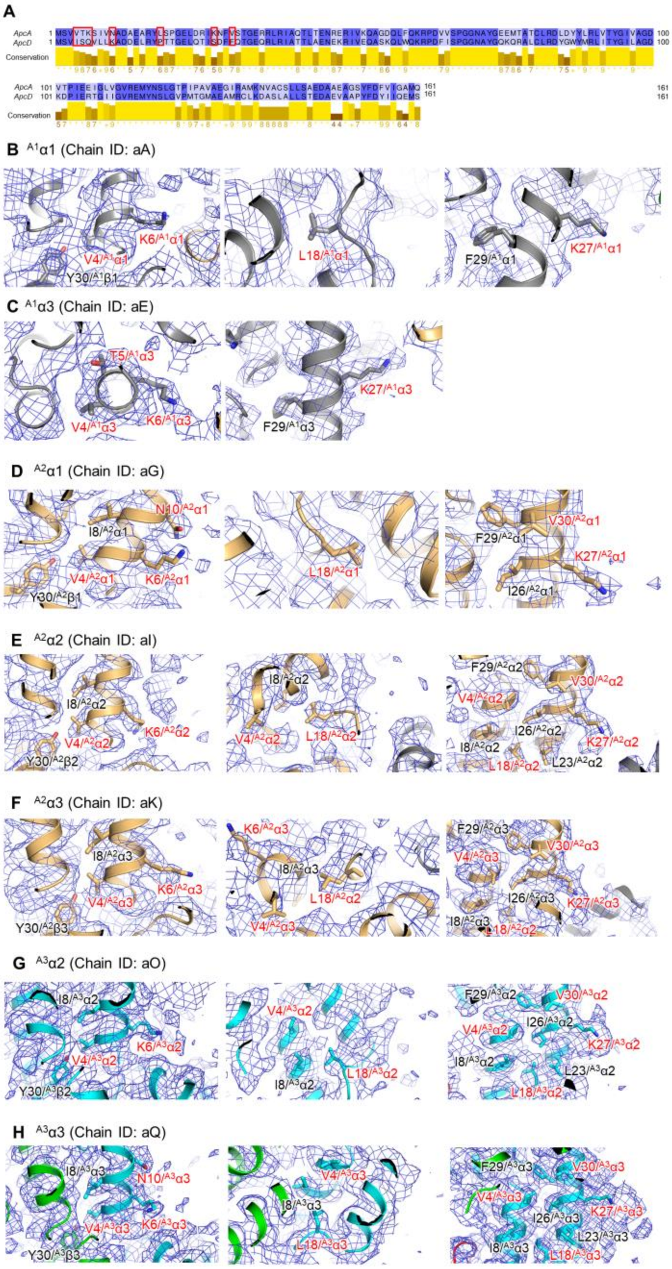


**Extended Data Figure 6. Identification of ApcD in the PBS core.** In this study, the subunit in the PBS core that could not be identified as ApcA (i.e., $^{A1}\alpha2$, Chain ID: aC/dC) was identified as ApcD. **(A)** Amino acid sequence alignment of ApcA and ApcD. The colors of the amino acid residues are represented in percentage identities using Jalview. The amino acid residues in the red boxes are residues that differ between ApcA and AcpD. **(B-H)** Each cryo-EM map in α subunit in the PBS core is shown in a mesh representation at 1.0–3.0 sigma contour levels.

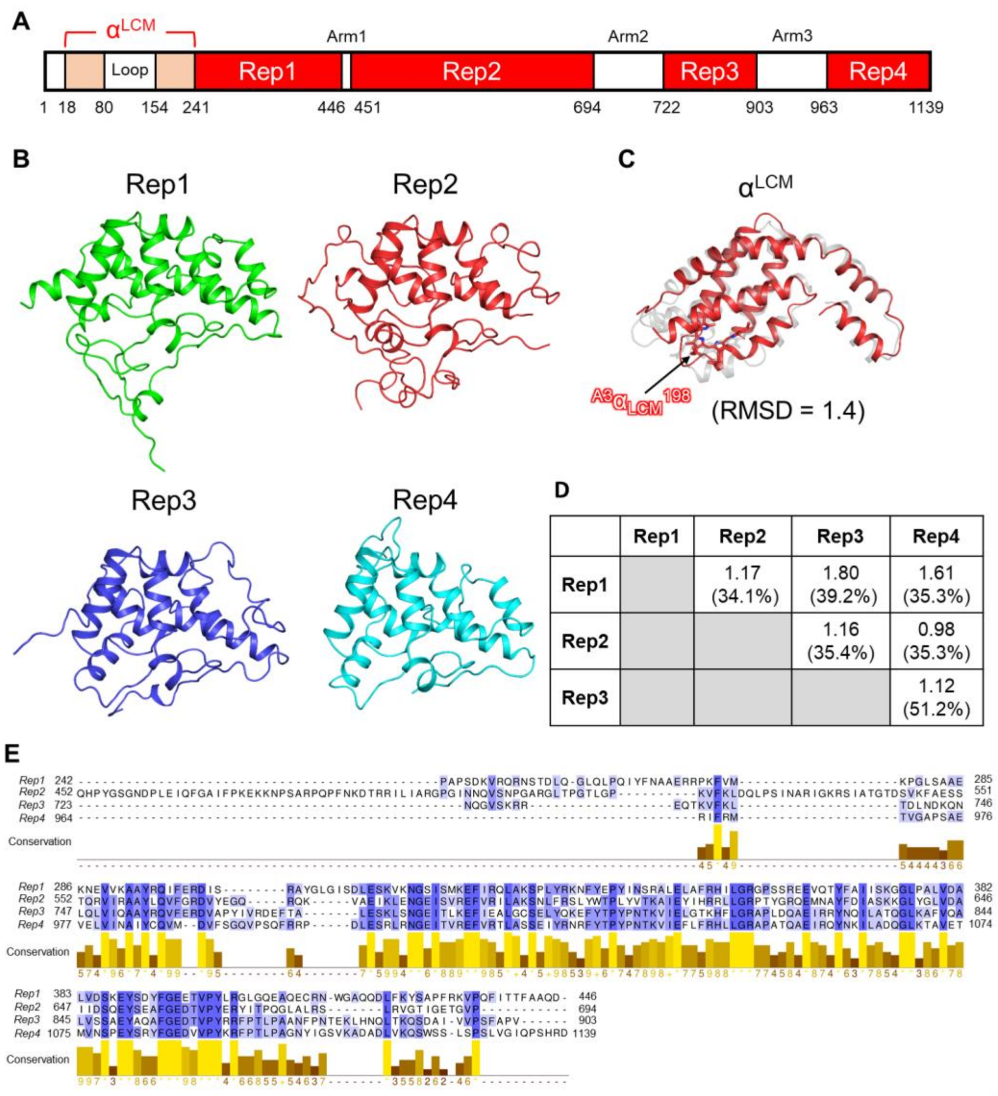


**Extended Data Figure 7. Structure of $\alpha^{LCM}$, Rep1–4 of ApcE ($L_{CM}$). (A)** Diagram of the structural element of ApcE. **(B)** Structures of Reg1–4. Reg1, green; Reg2, red; Reg3, blue; Reg4, cyan. **(C)** Superposition with $\alpha^{LCM}$ and ApcA. $\alpha^{LCM}$, red; transparent gray, crystal structure of APC (PDB code: 3DBJ). The value in parenthesis indicates the root mean square deviations (RMSD). Secondary structure matching in CCP4 was used to calculate the value of RMSD. **(D)** The values indicate RMSD. The values in parentheses indicate sequence identities. (E) Amino acid sequence alignment of the

Rep1–4 of $L_{CM}$. The colors of the amino acid residues are indicated by percentage identities using Jalview.

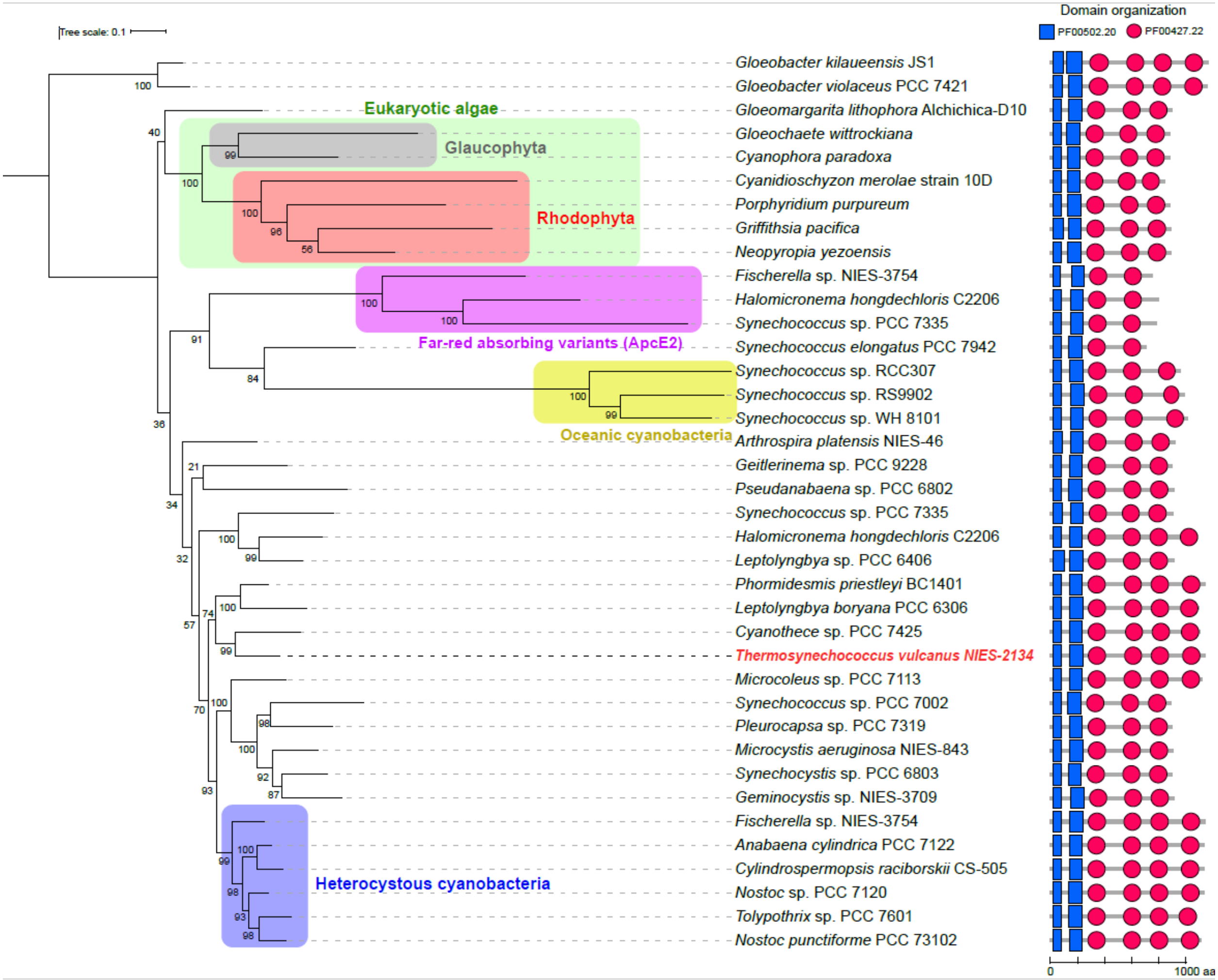


**Extended Data Figure 8. Phylogenic analysis of ApcE ($L_{CM}$) subunit.**

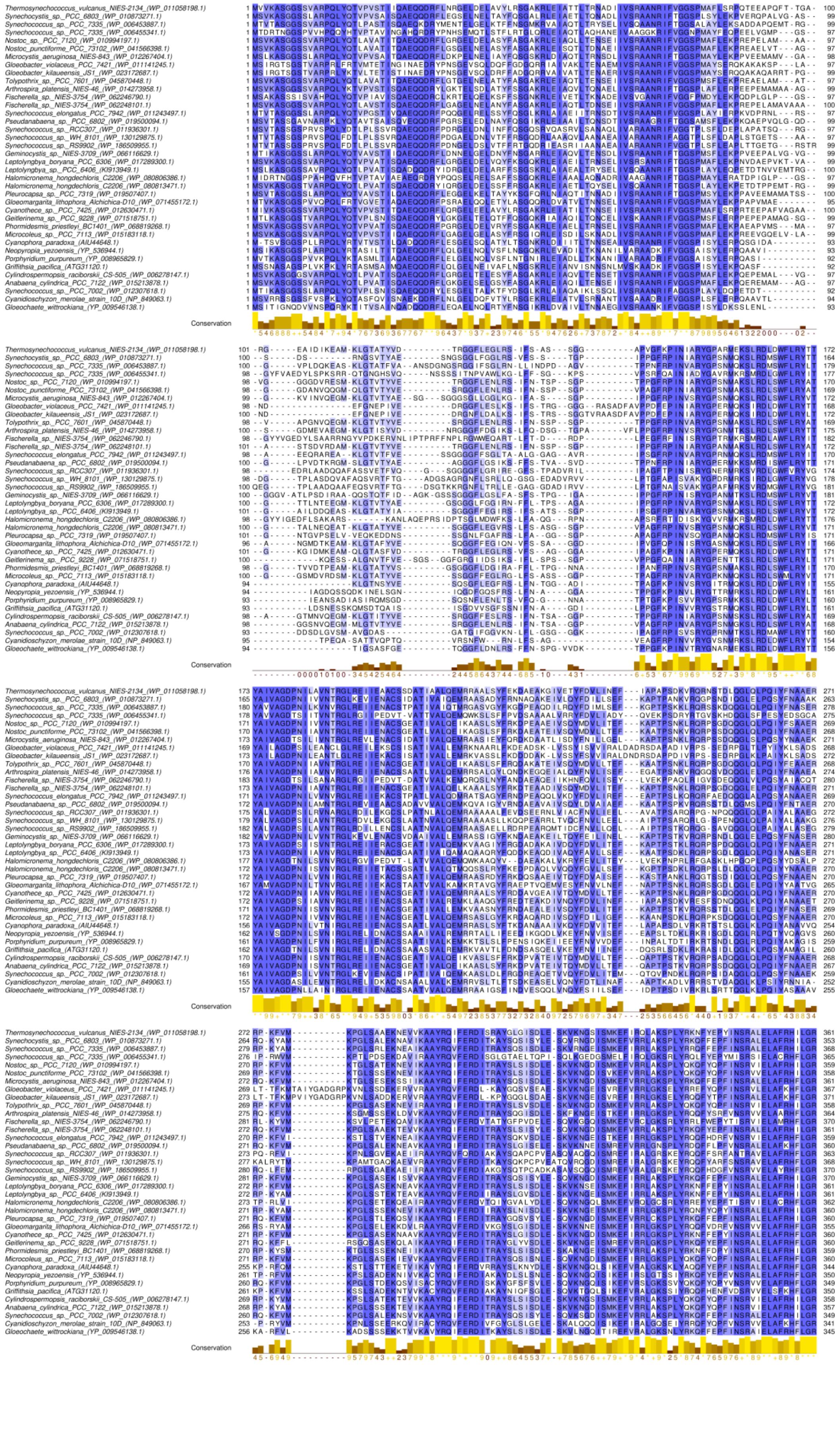

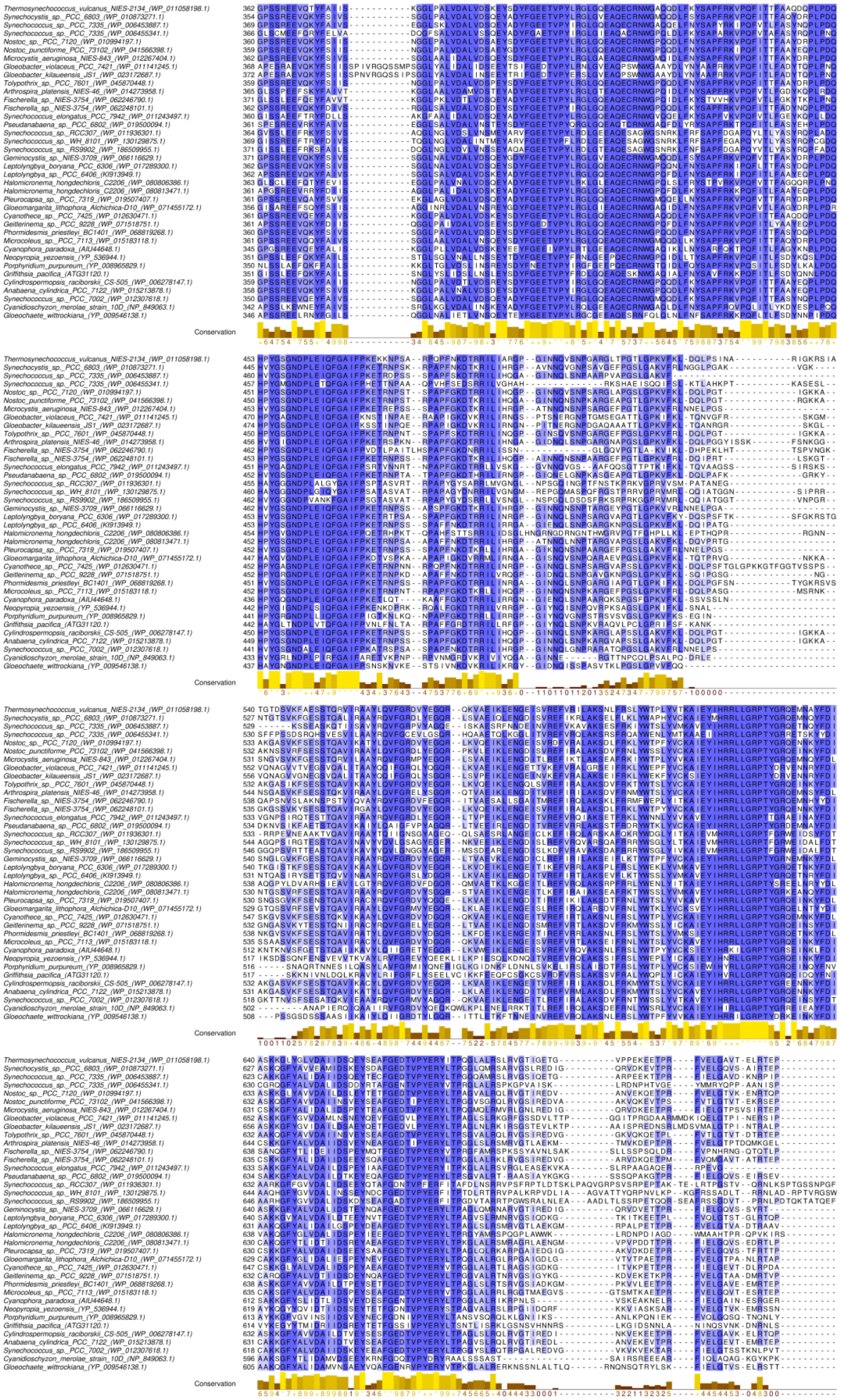
Conservation

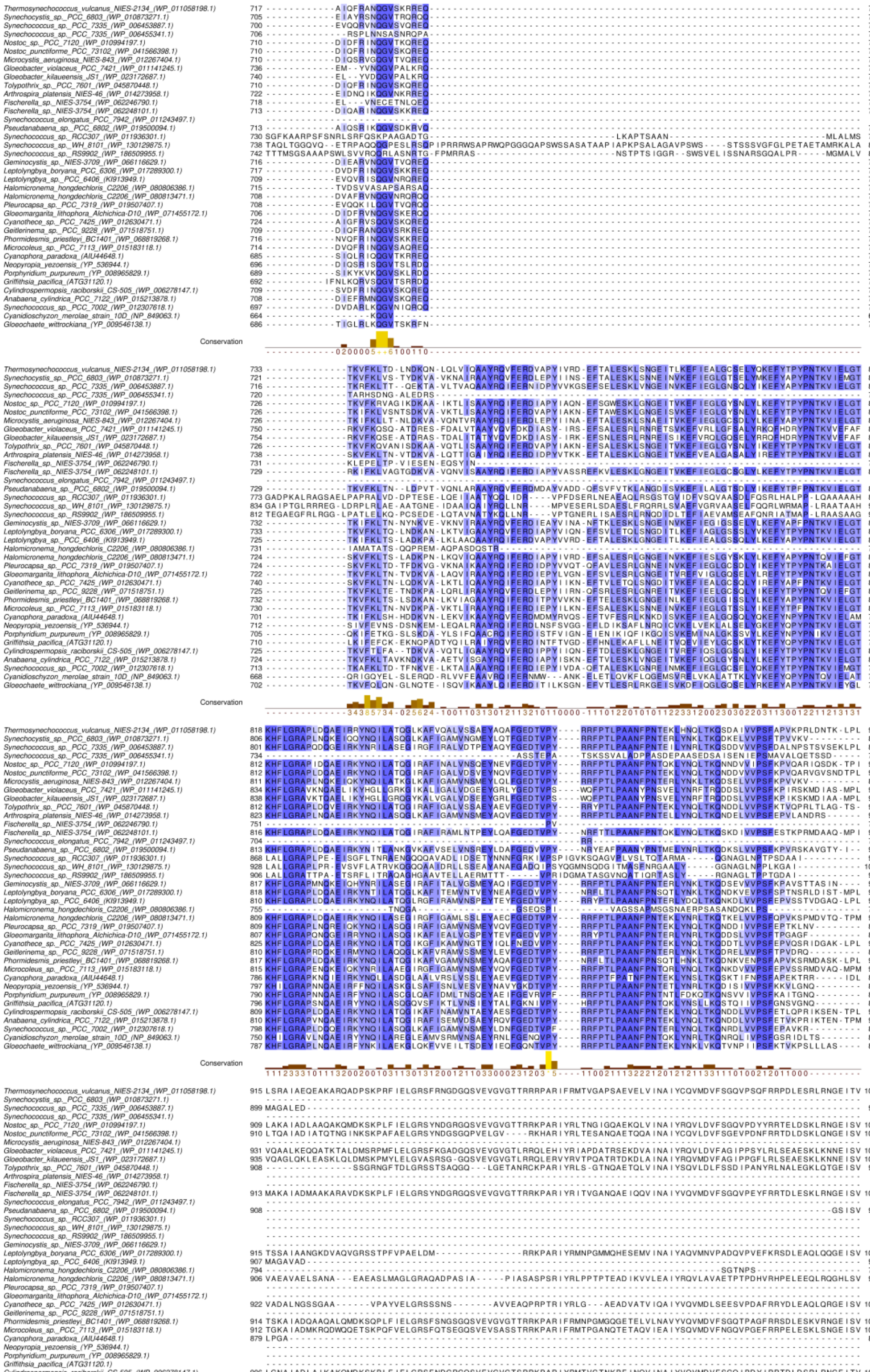

Conservation

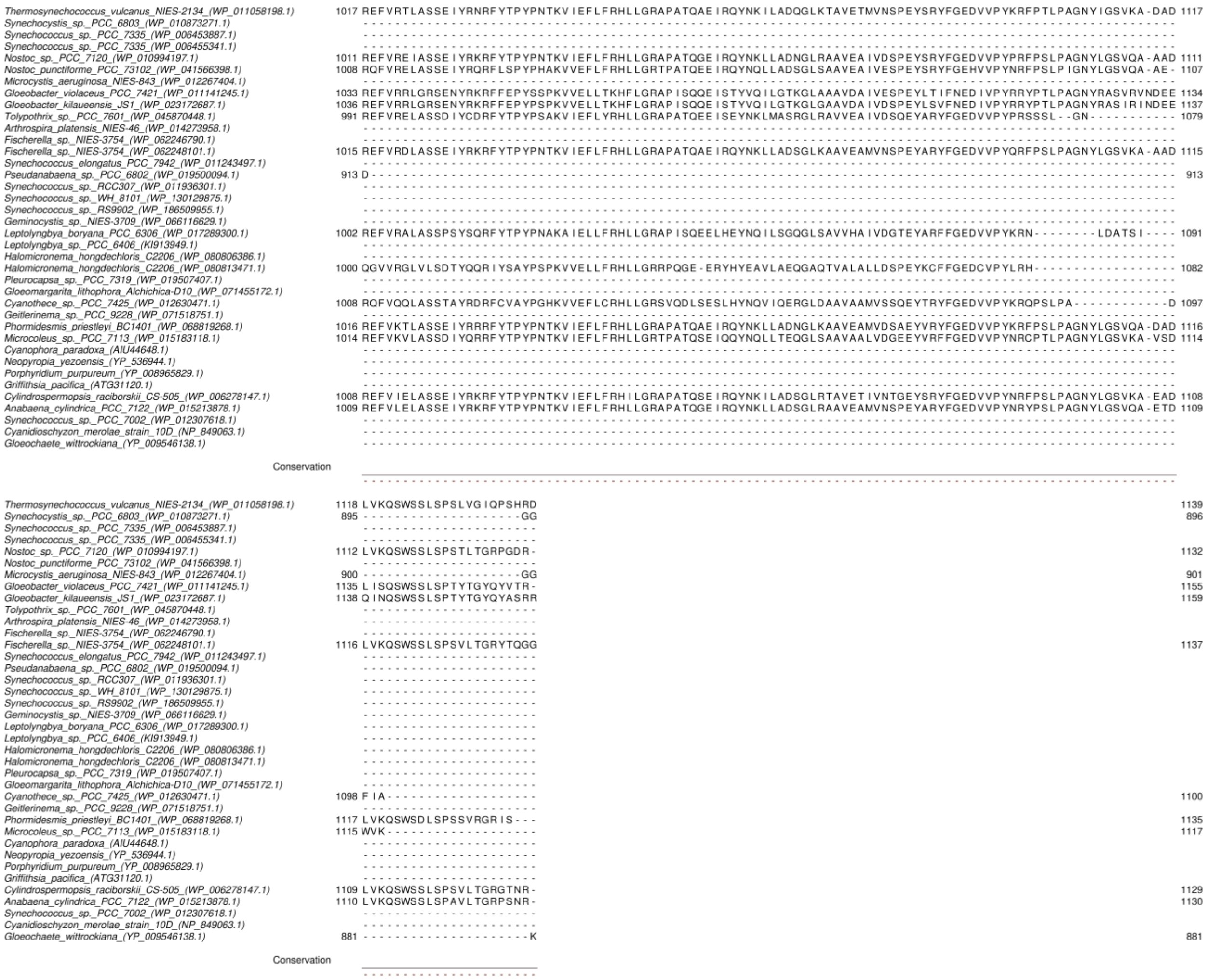

**Extended Data Figure 9. Amino acid sequence alignment of ApcE ($L_{CM}$) subunit from selected strains of cyanobacteria, glaucophytes, and rhodophytes.** The colors of the amino acid residues are indicated by percentage identities using Jalview.

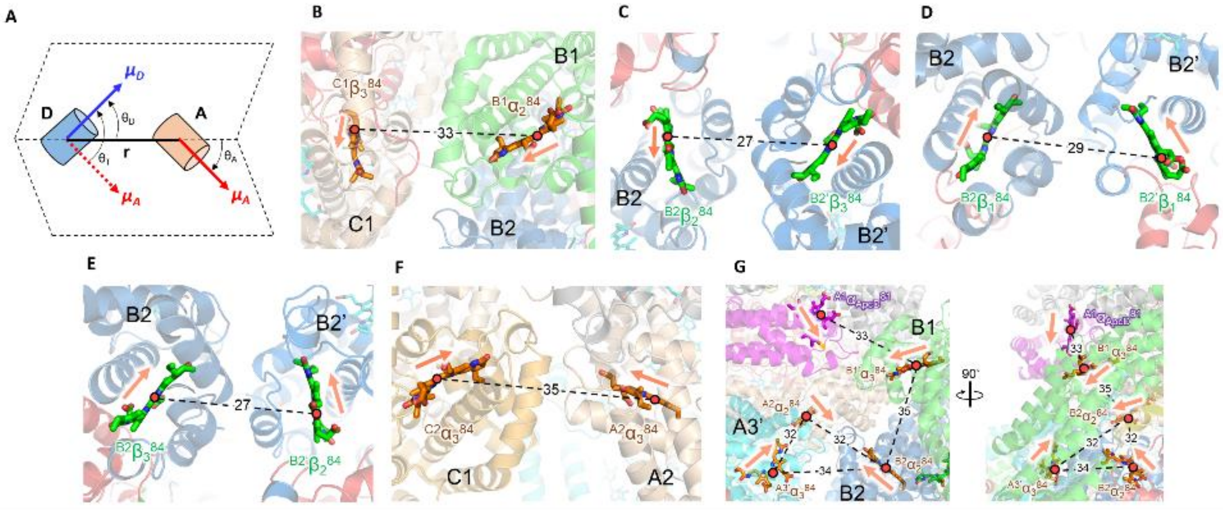


**Extended Data Figure 10. Arrangement of chromophores between cylinders. (A)** Schematic model of the Förster resonance energy transfer. The transition dipole moment of the donor (D) and acceptor (A) are $\boldsymbol{\mu_D}$ and $\boldsymbol{\mu_A}$, respectively. **r** is the intermolecular center-to-center distance between D and A. $\theta_D$ is the angle between D–A connecting line and D transition dipole moment. $\theta_A$ is the angle between D–A connecting line and A transition dipole moments. $\theta_T$ is the angle between D and A transition dipole moment. **(B-G)** Chromophores in the B and C cylinders that may be associated with the energy transfer. The numbers on the dotted lines indicate the distances (Å) between the PCB pairs. The orange arrows indicate the direction of the transition dipole moment for each chromophore. The orientation factor, □$^2$, is estimated using the following formula: $\kappa^2 = (\cos\theta_T - 3\cos\theta_D\cos\theta_A)^2$.

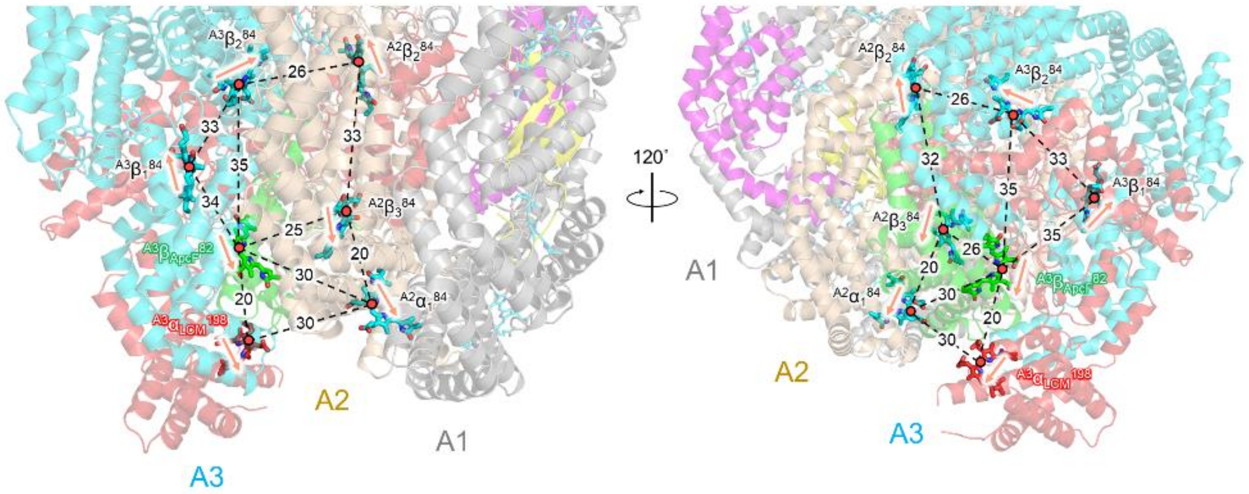


**Extended Data Figure 11. Arrangement of chromophores around the terminal emitters.** Chromophores in the A cylinders that may be associated with the energy transfer to the terminal emitters. The numbers on the dotted lines indicate the distances (Å) between the PCB pairs. The orange arrows indicate the direction of the transition dipole moment for each chromophore.

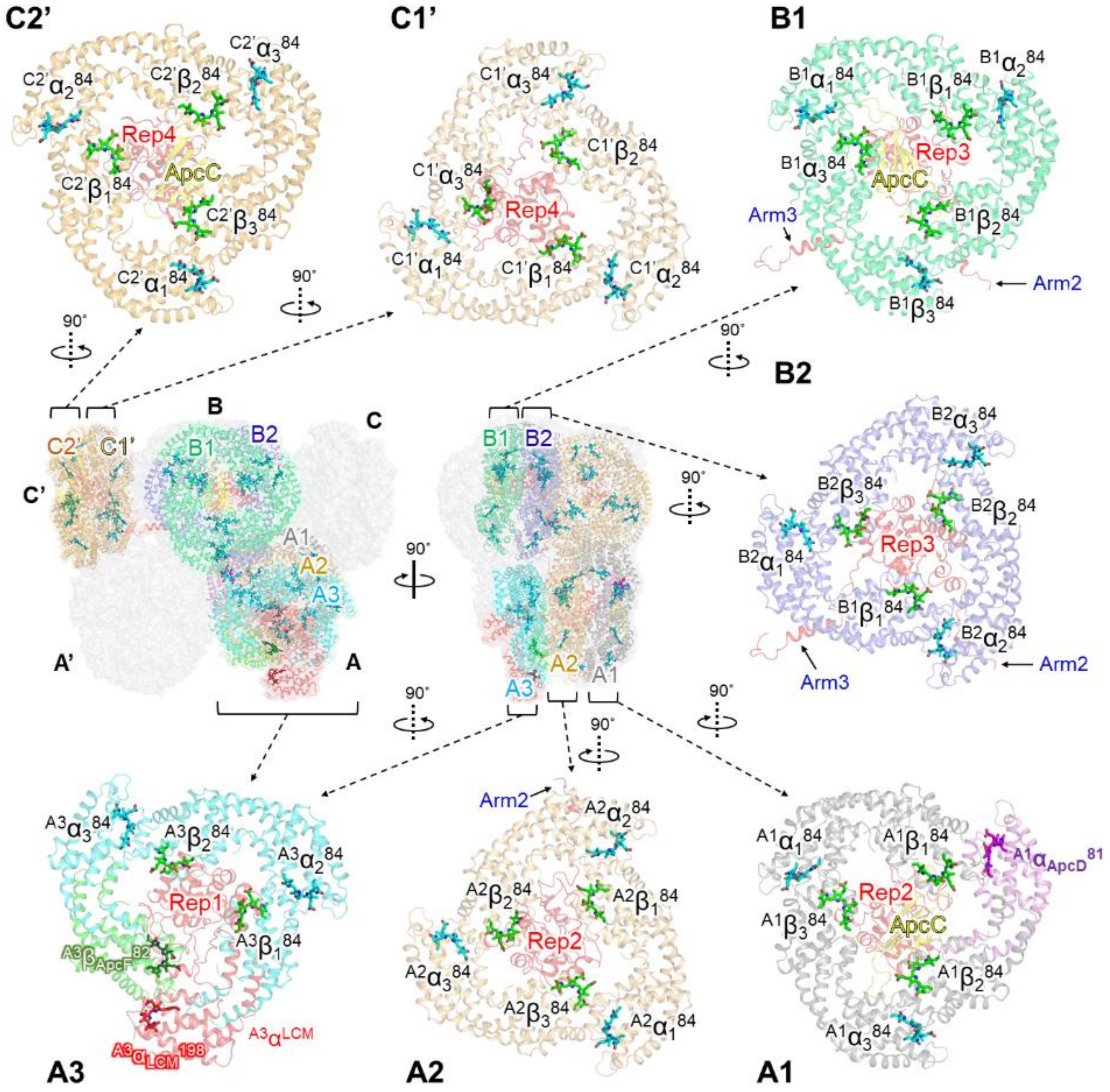


**Extended Data Figure 12. Arrangement of chromophores in the PBS core and their names.** PCBs bound in ApcA, cyan; PCBs bound in ApcB, green; PCBs bound in terminal emitters (ApcD, ApcE, and ApcF), magenta, red, and dark green, respectively.

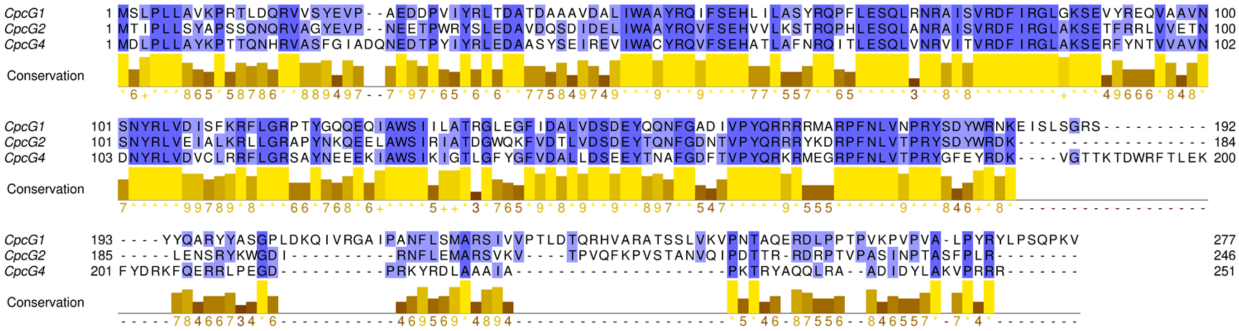


**Extended Data Figure 13. Amino acid sequence alignment of CpcG1, CpcG2, and CpcG4 of *T. vulcanus*.** The colors of the amino acid residues are indicated by percentage identities using Jalview.

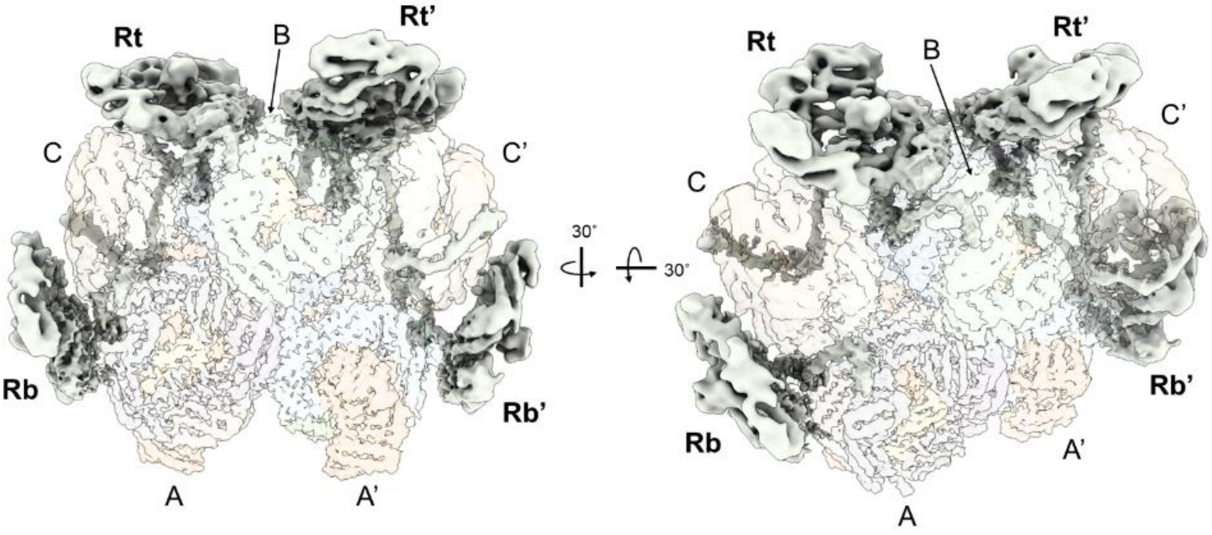


**Extended Data Figure 14. The PC rods interacting with PBS core from *T. vulcanus*.** The linker proteins extending from the PC rods (Rt, Rt’, Rb, and Rb’) interact with the A, B, and C cylinders of the PBS core. The PC rods and PBS core are shown in gray and translucent, respectively.

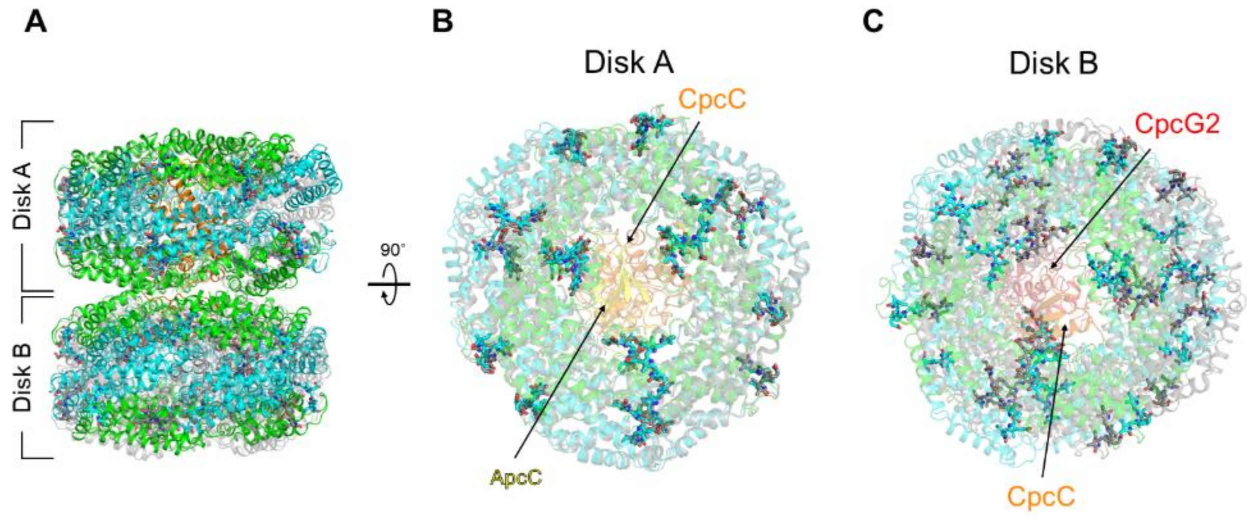


**Extended Figure 15. The structure of the PC rod as determined by cryo-EM and X-ray crystal structure analysis. (A)** Superposition with the PC rod visualized by cryo-EM and the X-ray crystal structure of PC. The PC rod by cryo-EM, multicolored; the crystal structure of PC (PDB code: 3O18), gray. **(B)** Structure of the PC rod rotated 90̊. The arrangement of chromophores in Disk A is shown. PCBs identified by cryo-EM, cyan; PCBs identified by X-ray crystal structure analysis, gray. **(C)** Structure of the PC rod rotated 90̊. The arrangement of chromophores in Disk B is shown. Colors are the same as in (B).

**Extended Data Table 1. Statistics for data collection, processing, and refinement.**

| **Data collection** | PBS core (PDB: 7VEA, EMDB-31944) | PC rod (PDB: 7VEB, EMDB-31945) |
|---|---|---|
| Microscope | CRYO ARM 300 | |
| Imaging device | K2 summit | |
| Accelerating voltage (kV) | 300 | |
| Imaging mode | Counting | |
| Data collection | JADAS | |
| Grid condition | Quantifoil R1.2/1.3 Cu 200 mesh Au sputtered | |
| Number of optics group | 8 | None |
| Nominal magnification | x40,000 | |
| Total exposure time (sec) | 6 | |
| No. of frames | 30 | 50 |
| Total electron exposure ($e^-/Å^2$) | 84.1 | 50.5 |
| Defocus range (µm) | -0.5 – -1.5 | |
| Original pixel size | 1.24 | |
| No. of total image sets | 4,600 | 2,865 |
| **Data processing** | | |
| No. of used image sets | 3,196 | 2,573 |
| Initial particles (no.) | 128,676 | 812,327 |
| Final particles (no.) | 25,532 | 159,537 |
| Pixel size for final map | 1.24 | 1.86 |
| Symmetry imposed | C2 | C1 |
| Map resolution (Å) | 3.72 | 4.19 |
| B-factor for sharpening | -39.6 | -89.3 |
| FSC threshold | 0.143 | 0.143 |
| Map resolution range ($Å^2$) | 3.28 – 20.73 | 4.02 – 6.96 |
| **Refinement** | | |
| Initial models (PDB code) | 3O18, 3DBJ, 5Y6P, 6KGX | |
| Model resolution (Å) | 3.82 | 4.24 |
| FSC threshold | 0.5 | |
| Model composition | | |
| Non-hydrogen atoms | 111,828 | 35,543 |
| Protein residues | 15,594 | 4,536 |

| | | |
|---|---|---|
| Ligands | 84 | 36 |
| B factors ($Å^2$) | | |
| Protein | 124.9 | 119.5 |
| Ligands | 126.1 | 121.5 |
| R.m.s. deviations | | |
| Bond lengths (Å) | 0.005 | 0.005 |
| Bond angles (Å) | 1.043 | 0.904 |
| Validation | | |
| MolProbity score | 1.37 | 1.51 |
| Clashscore | 6.69 | 6.62 |
| Poor rotamers (%) | 0.66 | 0.68 |
| Ramachandran plot | | |
| Favored (%) | 98.2 | 97.2 |
| Allowed (%) | 1.7 | 2.7 |
| Disallowed (%) | 0.1 | 0.1 |

**Extended Data Table 2. Evaluation of the agreement between cryo-EM map and final model (PBS core).**

| Chain ID | Protein | Q-score* | Estimated resolution (Å) | Chain ID | Protein | Q-score* | Estimated resolution (Å) |
|---|---|---|---|---|---|---|---|
| aA, dA | ApcA | 0.52 | 3.37 | bQ, eQ | ApcA | 0.48 | 3.62 |
| aB, dB | ApcB | 0.40 | 4.04 | bR, eR | ApcB | 0.53 | 3.34 |
| aC, dC | ApcD | 0.39 | 4.09 | bS, eS | ApcA | 0.40 | 4.05 |
| aD, dD | ApcB | 0.37 | 4.20 | bT, eT | ApcB | 0.46 | 3.71 |
| aE, dE | ApcA | 0.42 | 3.91 | bU, eU | ApcA | 0.49 | 3.52 |
| aF, dF | ApcB | 0.43 | 3.91 | bV, eV | ApcB | 0.43 | 3.86 |
| aG, dG | ApcA | 0.54 | 3.29 | bW, eW | ApcA | 0.46 | 3.72 |
| aH, dH | ApcB | 0.61 | 2.85 | bX, eX | ApcB | 0.45 | 3.78 |
| aI, DI | ApcA | 0.61 | 2.87 | bY, eY | ApcC | 0.55 | 3.20 |
| aJ, dJ | ApcB | 0.59 | 2.97 | cA, fA | ApcA | 0.41 | 3.98 |
| aK, dK | ApcA | 0.54 | 3.26 | cB, fB | ApcB | 0.41 | 3.98 |
| aL, dL | ApcB | 0.56 | 3.12 | cC, fC | ApcA | 0.40 | 4.03 |
| aM, dM | ApcE | 0.56 | 3.13 | cD, fD | ApcB | 0.51 | 3.42 |
| aN, dN | ApcB | 0.52 | 3.35 | cE, fE | ApcA | 0.50 | 3.51 |
| aO, dO | ApcA | 0.51 | 3.45 | cF, fF | ApcB | 0.49 | 3.52 |
| aP, dP | ApcB | 0.60 | 2.94 | cG, fG | ApcA | 0.36 | 4.30 |
| aQ, dQ | ApcA | 0.59 | 2.96 | cH, fH | ApcB | 0.37 | 4.22 |
| aR, dR | ApcF | 0.56 | 3.15 | cI, fI | ApcA | 0.40 | 4.06 |
| aS, dS | ApcC | 0.43 | 3.87 | cJ, fJ | ApcB | 0.44 | 3.81 |
| bM, eM | ApcA | 0.52 | 3.35 | cK, fK | ApcA | 0.48 | 3.61 |
| bN, eN | ApcB | 0.59 | 2.96 | cL, fL | ApcB | 0.39 | 4.08 |
| bO, eO | ApcA | 0.59 | 3.00 | cM, fM | ApcC | 0.51 | 3.46 |
| bP, eP | ApcB | 0.56 | 3.13 | Total | | 0.49 | 3.55 |

*Q-score: An index showing resolvability of atoms, amino acid residues, and ligands assigned in a cryo-EM map (local resolution map). Resolution (Å) was estimated using the formula (Q-score = −0.1775× Resolution (Å) + 1.1192) according to Pintilie et al. (2020).

**Extended Data Table 3. Evaluation of the agreement between cryo-EM map and final model (PC rod).**

| Chain ID | Protein | Q-score* | Estimated resolution (Å) |
|---|---|---|---|
| A | CpcA | 0.42 | 3.95 |
| B | CpcB | 0.41 | 3.98 |
| C | CpcA | 0.39 | 4.12 |
| D | CpcB | 0.41 | 3.97 |
| E | CpcA | 0.42 | 3.91 |
| F | CpcB | 0.44 | 3.84 |
| G | CpcA | 0.42 | 3.93 |
| H | CpcB | 0.42 | 3.91 |
| I | CpcA | 0.40 | 4.04 |
| J | CpcB | 0.41 | 3.98 |
| K | CpcA | 0.41 | 3.99 |
| L | CpcB | 0.44 | 3.83 |
| M | CpcA | 0.42 | 3.97 |
| N | CpcB | 0.43 | 3.90 |
| O | CpcA | 0.36 | 4.26 |
| P | CpcB | 0.43 | 3.90 |
| Q | CpcA | 0.33 | 4.42 |
| R | CpcB | 0.36 | 4.30 |
| S | CpcA | 0.38 | 4.14 |
| T | CpcB | 0.33 | 4.42 |
| U | CpcA | 0.40 | 4.06 |
| V | CpcB | 0.42 | 3.94 |
| W | CpcA | 0.31 | 4.57 |
| X | CpcB | 0.35 | 4.33 |
| Y | CpcD | 0.48 | 3.60 |
| Z | CpcC | 0.47 | 3.64 |
| a | CpcG2 | 0.41 | 4.01 |
| Total | | 0.40 | 4.03 |

*Q-score: An index showing resolvability of atoms, amino acid residues, and ligands assigned in a cryo-EM map (local resolution map). Resolution (Å) was estimated using the formula (Q-score = −0.1775× Resolution (Å) + 1.1192) according to Pintilie et al. (2020).

**Extended Data Table4. Subunit protein in PBS core and PC rod of *T. vulcanus*.**

| Structure (PDB code) | Name | Protein | M. W. /kDa* | Number of molecules in a structural model | Remarks |
|---|---|---|---|---|---|
| PBS core (7VEA) | α subunit | ApcA | 17.5 | 38 | Main components of the PBS core |
| | β subunit | ApcB | 17.4 | 40 | Main components of the PBS core |
| | Core linker ($L_C$) | ApcC | 7.9 | 6 | Core-cap linker |
| | α subunit | ApcD | 18.1 | 2 | Terminal emitter |
| | Core-membrane linker ($L_{CM}$) | ApcE | 127.6 | 2 | Terminal emitter. Components in the subunit: α, Reps 1–4, and Arms 1–3. |
| | β subunit | ApcF | 18.7 | 2 | Terminal emitter |
| PC rod (7VEB) | α subunit | CpcA | 17.4 | 12 | Main components of PC rod |
| | β subunit | CpcB | 18.2 | 12 | Main components of PC rod |
| | Rod linker ($L_R$) | CpcC | 32.1 | 1 | Phycocyanin-associated rod linker |
| | Rod-terminal linker ($L_{RT}$) | CpcD | 8.7 | 1 | Rod-cap linker |
| | Rod-core linker ($L_{RC}$) | CpcG1 | 31.6 | 0 | Phycocyanin-associated rod-core linker |
| | Rod-core linker ($L_{RC}$) | CpcG2 | 28.8 | 1 | Phycocyanin-associated rod-core linker |
| | Rod-core linker ($L_{RC}$) | CpcG4 | 29.6 | 0 | Phycocyanin-associated rod-core linker |

*The molecular weight (M. W.) of each subunit is estimated from the composition of the amino acid residues in each subunit.

**Extended Data Table 5. Estimation of the orientation factor between major chromophores in the PBS core.**

| Pigment (D or A) | Pigment (A or D) | Orientation factor ($\kappa^2$) | Distance (Å) |
|---|---|---|---|
| $^{C1}\beta_3^{84}$ | $^{B1}\alpha_2^{84}$ | 1.0 | 33 |
| $^{B2}\beta_2^{84}$ | $^{B2'}\beta_3^{84}$ | 0.8 | 27 |
| $^{B2}\beta_1^{84}$ | $^{B2'}\beta_1^{84}$ | 1.0 | 29 |
| $^{B2}\beta_3^{84}$ | $^{B2'}\beta_2^{84}$ | 0.8 | 27 |
| $^{C2}\alpha_3^{84}$ | $^{A2}\alpha_3^{84}$ | 3.0 | 35 |
| $^{A1}\alpha_{ApcD}^{81}$ | $^{B1'}\alpha_3^{84}$ | 3.0 | 33 |
| $^{B1'}\alpha_3^{84}$ | $^{B2}\alpha_2^{84}$ | 1.6 | 35 |
| $^{B2}\alpha_2^{84}$ | $^{A2}\alpha_2^{84}$ | 2.9 | 32 |
| $^{B2}\alpha_2^{84}$ | $^{A3'}\alpha_3^{84}$ | 2.5 | 34 |
| $^{A2}\alpha_2^{84}$ | $^{A3'}\alpha_3^{84}$ | 2.1 | 32 |
| $^{A3}\alpha_{LCM}^{198}$ | $^{A3}\beta_{ApcF}^{82}$ | 2.9 | 20 |
| $^{A3}\alpha_{LCM}^{198}$ | $^{A2}\alpha_1^{84}$ | 1.9 | 30 |
| $^{A3}\beta_{ApcF}^{82}$ | $^{A2}\alpha_1^{84}$ | 1.6 | 30 |
| $^{A3}\beta_{ApcF}^{82}$ | $^{A2}\beta_3^{84}$ | 1.3 | 25 |
| $^{A3}\beta_{ApcF}^{82}$ | $^{A3}\beta_2^{84}$ | 0.9 | 35 |
| $^{A3}\beta_{ApcF}^{82}$ | $^{A3}\beta_1^{84}$ | 1.9 | 34 |
| $^{A2}\alpha_1^{84}$ | $^{A2}\beta_3^{84}$ | 2.3 | 20 |
| $^{A3}\beta_2^{84}$ | $^{A2}\beta_3^{84}$ | 1.2 | 33 |
| $^{A2}\beta_2^{84}$ | $^{A3}\beta_2^{84}$ | 1.1 | 26 |
| $^{A3}\beta_1^{84}$ | $^{A3}\beta_2^{84}$ | 1.3 | 33 |

The distance (Å) between a pair of chromophores (donor (D) and acceptor (A)) and their orientation factor ($\kappa^2$).
